# ChatNT: A Multimodal Conversational Agent for DNA, RNA and Protein Tasks

**DOI:** 10.1101/2024.04.30.591835

**Authors:** Guillaume Richard, Bernardo P. de Almeida, Hugo Dalla-Torre, Christopher Blum, Lorenz Hexemer, Priyanka Pandey, Stefan Laurent, Marie Lopez, Alexandre Laterre, Maren Lang, Uğur Şahin, Karim Beguir, Thomas Pierrot

**Author notes:** Equal contributions (Guillaume Richard, Bernardo P. de Almeida). Corresponding author(s) © 2024 InstaDeep. All rights reserved.

## Abstract

Language models are thriving, powering conversational agents that assist and empower humans to solve a number of tasks. Recently, these models were extended to support additional modalities including vision, audio and video, demonstrating impressive capabilities across multiple domains including healthcare. Still, conversational agents remain limited in biology as they cannot yet fully comprehend biological sequences. On the other hand, high-performance foundation models for biological sequences have been built through self-supervision over sequencing data, but these need to be fine-tuned for each specific application, preventing transfer and generalization between tasks. In addition, these models are not conversational which limits their utility to users with coding capabilities. In this paper, we propose to bridge the gap between biology foundation models and conversational agents by introducing ChatNT, the first multimodal conversational agent with an advanced understanding of biological sequences. ChatNT achieves new state-of-the-art results on the Nucleotide Transformer benchmark while being able to solve all tasks at once, in English, and to generalize to unseen questions. In addition, we have curated a new set of more biologically relevant instructions tasks from DNA, RNA and proteins, spanning multiple species, tissues and biological processes. ChatNT reaches performance on par with state-of-the-art specialized methods on those tasks. We also present a novel perplexity-based technique to help calibrate the confidence of our model predictions. Our framework for genomics instruction-tuning can be easily extended to more tasks and biological data modalities (e.g. structure, imaging), making it a widely applicable tool for biology. ChatNT is the first model of its kind and constitutes an initial step towards building generally capable agents that understand biology from first principles while being accessible to users with no coding background.

## Introduction

Understanding how cells, tissues, and organisms interpret information encoded in the genome is of paramount importance for advancing our comprehension of biology. The DNA sequence of an organism comprises all the instructions to specify RNAs and proteins, but also when and in which cellular context these should be produced. Since the human genome was sequenced [1], the main focus has been on identifying every genomic element, characterizing their function, and assessing the impact of genetic variants on the different gene regulatory and cellular processes. Given the complexity of biological sequences and processes, and the increasing volume of genomics data, several machine learning and deep learning methods have been developed to address these questions by predicting diverse molecular phenotypes with great accuracy [2–4]. These tasks include predicting the binding of proteins to DNA and RNA [5, 6], DNA methylation [7], chromatin features [8–10], regulatory elements [11], 3D genome folding [12–14], splicing [15, 16], gene expression [10, 17, 18], mRNA properties such as stability [19] and polyadenylation [20, 21], and protein properties such as melting point [22].

While supervised deep learning models have already significantly improved the predictive capabilities on these tasks, their performance remains often limited due to the scarcity of labeled data, given that labelling is time consuming and expensive. On the other hand, an exponentially increasing volume of raw genome data is becoming available thanks to the increase in throughput and reduced cost of modern sequencing techniques, thus creating a significant opportunity for self-supervised deep learning methods to train on such unlabeled data. Through learning-techniques such as masked-or next-token prediction [23–25], with tokens representing one or several consecutive nucleotides, deep learning models can build powerful foundation representations of the genome during this “pre-training” stage, aggregating correlations between nucleotides and larger sequence patterns into rich high-dimensional vectors that capture known genomic elements and protein binding sites [26]. These models can later exploit these rich representations, during a “fine-tuning” stage, to learn faster and reach better performance on supervised tasks, i.e. tasks where labels are available, despite data scarcity. Recently, several such foundation models have been built in this fashion, showing that they can be pre-trained on the genomes of hundreds of species before being fine-tuned to solve a large collection of molecular phenotype prediction tasks [26–32].

This being said, the performance and application domain of current DNA foundation models remains limited. In the current paradigm, foundation models require fine-tuning to each specific task individually to produce accurate representations and predictions, and are thus better characterized as narrow experts on specific tasks. This not only yields a deluge of different models as the number of tasks increases, but also prevents any transfer between supervised tasks as well as to solve new tasks in a zero-shot setting (i.e. without the need for further finetuning on some examples). There is therefore a need to rethink the development of genomics Artificial Intelligence (AI) systems with the goal of establishing general, unified models that capture the intricate relationships between all diverse biological sequences and functions. It has been shown in other fields such as natural language processing (NLP) and computer vision that training on several tasks in parallel results in knowledge transfer between tasks and improved accuracy and generalization [23, 24, 33–35]. In these domains, English language has been shown to play a wider role: a universal interface for representing various tasks and instructions and helping guide the training of end-to-end multi-task models [36, 37]. Transferring this type of approaches to biological data is a promising approach towards developing a general model that can solve all genomics tasks of interest simultaneously and with improved accuracy.

An additional important aspect of building a universal genomics AI system is its accessibility to different types of users. Most biologists do not know how to use current genomics models, let alone how to program one themselves for a given task of interest. Such models are not conversational and thus of limited utility in practice to users with no coding capabilities. Also here, language can play an important role as a universal interface for a general-purpose AI assistant that can solve genomics tasks through task instructions that can be explicitly represented in English language. For example, the recent success of ChatGPT [38] and GPT-4 [39] has demonstrated the power of large language models (LLMs) trained to follow human instructions, and how such tools can transform several industries due to their ease of use. We envision the same paradigm shift for genomics and biology once we have “ChatGPT-like” agents that are proficient in biological tasks.

To that end, we introduce in this work a novel approach to build foundation models for genomics. Similarly to lines of works that emerged in NLP [24, 25, 36], and inspired by recent vision/language multimodal models [40–46], we propose to formulate all supervised genomics prediction tasks as text-to-text tasks and to build a multi-modal DNA/language agent, dubbed the Chat Nucleotide Transformer (or ChatNT). ChatNT can be given one or several DNA sequences and is prompted in English to solve all those tasks. This formulation allows us to express all tasks with the same vocabulary, being here the concatenation of the English and DNA vocabularies, and to learn to solve them by minimizing a unified objective, similar to GPT-like models [25, 47], allowing for seamless new task integration and generalization. Formulating tasks in English is also an easy way to provide additional meta-data information to the model, such as the species, the chromosome or the cell type, that is also missing in most current DNA foundation models.

ChatNT is built to act as a generalist genomics AI system - a unified model that can interpret multiple biological sequences and handle dozens of tasks in a conversational agent setting. To the best of our knowledge, ChatNT is the first multimodal bio-sequence/English agent. We created the first datasets of genomics instructions tasks with curated sets of questions and instructions in English for diverse classification and regression tasks. We first show that ChatNT achieves a new state-of-the-art on the Nucleotide Transformer benchmark [26]. We next evaluate ChatNT in additional biologically relevant tasks that cover DNA, RNA and protein processes. ChatNT achieves state-of-the-art performance across all tasks, matching the performance of several specialized models, such as APARENT2 for RNA polyadenylation [20, 21] and ESM2 for protein-related tasks [48], while being able to solve a large collection of tasks at once and in English. Finally, its English conversational capabilities make its use easier than other models, widening its accessibility to scientists with no machine learning or computer science background. This framework for genomics instruction-tuning can be easily extended to new tasks or biological data modalities (e.g. sequencing experiments, imaging) without the need for pre-training from scratch every time, making it a widely applicable tool for biology.

## Results

### ChatNT: a unified framework to transform DNA foundation models into conversational agents to solve multiple tasks

ChatNT is the first framework for genomics instruction-tuning, extending instruction-tuning agents to the multimodal space of biology and biological sequences. Our framework is designed to be modular and trainable end-to-end. It combines (1) a DNA encoder model, pre-trained on raw genome sequencing data and that provides DNA sequence representations; (2) an English decoder, typically a pre-trained GPT-style LLM, to comprehend the user instructions and produce responses; and (3) a projection layer that projects the representations extracted by the DNA encoder into the embedding space of the input English words, such that both can be used by the English decoder (Fig. 1c; see Methods). In contrast to most multimodal works (e.g. [40, 49]) that would typically freeze the encoder and train only the projection, and sometimes the decoder, we decided in this work to backpropagate the gradients in the encoder in addition to the projection to allow supervised knowledge propagation at the DNA model level. The English decoder is kept frozen and therefore ChatNT benefits from its entire initial conversational capabilities, ensuring these do not degrade during training. In this work, we use the Nucleotide Transformer v2 (500M) model for the DNA encoder part [26] and Vicuna-7b (instruction-fine-tuned LLaMA model with 7B parameters) for the English decoder part [50] in order to build the conversational agent ChatNT. Keeping this modular architecture allows to use constantly improving encoders and decoders in the future without changing the model architecture.

**Figure 1.**
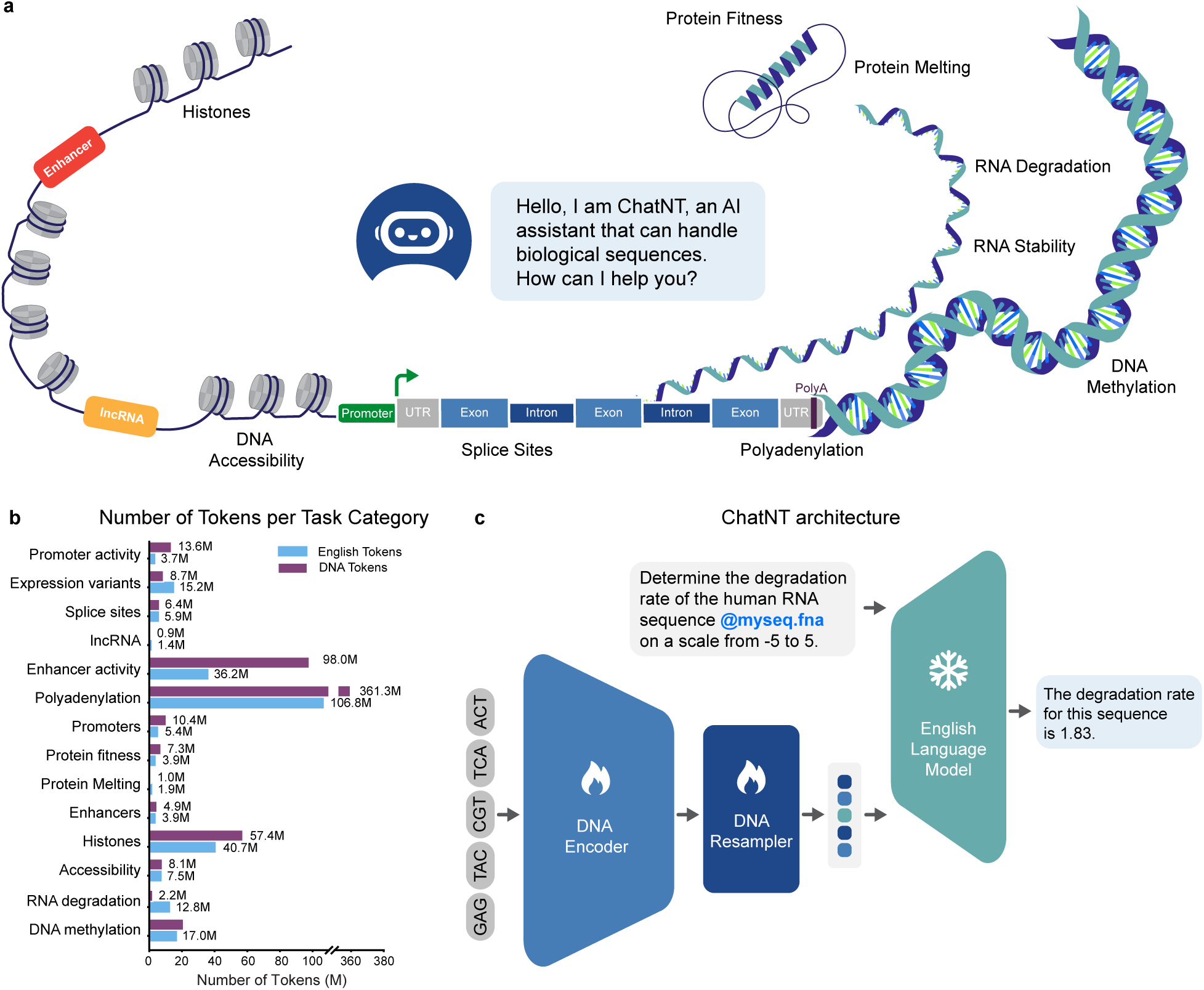
ChatNT, a conversational agent that can be prompted to solve a variety of biological tasks. **a.** Illustration of the different categories of downstream tasks included during training. **b.** Statistics about the number of English and DNA tokens available for each task in our genomics instructions dataset. English question/answer instructions are tokenized with the LLaMA tokenizer [47] while DNA sequences are tokenized using the Nucleotide Transformer tokenizer [26]. **c.** ChatNT approach to build a multimodal and multi-task genomics AI system. ChatNT conversational agent can be prompted in English to solve various tasks given an input question and nucleotide sequence. In this example, the user inputs a DNA sequence (fasta file) and asks the agent to evaluate the degradation rate of the given RNA sequence. The question tokens are combined with the projected DNA representations before passing through the English Language Model decoder. The pre-trained decoder writes the answer through next-token prediction, in this case predicting the degradation rate of the input sequence.

To train and evaluate ChatNT, we converted datasets of genomics tasks into instructions datasets by framing each task in English (Fig. 2; see Methods and respective results sections). We created for every task a train and test file each containing the respective DNA sequences combined with curated questions and answers in English. See Figure 1c for an example of question and answer for predicting RNA degradation levels: “*User: Determine the degradation rate of the human RNA sequence @myseq.fna on a scale from -5 to 5. ChatNT: The degradation rate for this sequence is 1.83.*”, where the projected embeddings of the candidate DNA sequence are inserted at the *@myseq.fna* position. We keep the same train/test splits as the original sources of each task, and use different questions for train and test to assess the English generalization capabilities of the model. This allows to not only evaluate the agent capability to generalize between DNA sequences but also its robustness to the English language used. We also provide a novel and flexible way to interleave English and DNA sequences through the usage of positional tags (*@myseq.fna*), allowing users to refer to several sequences in the same question.

**Figure 2.**
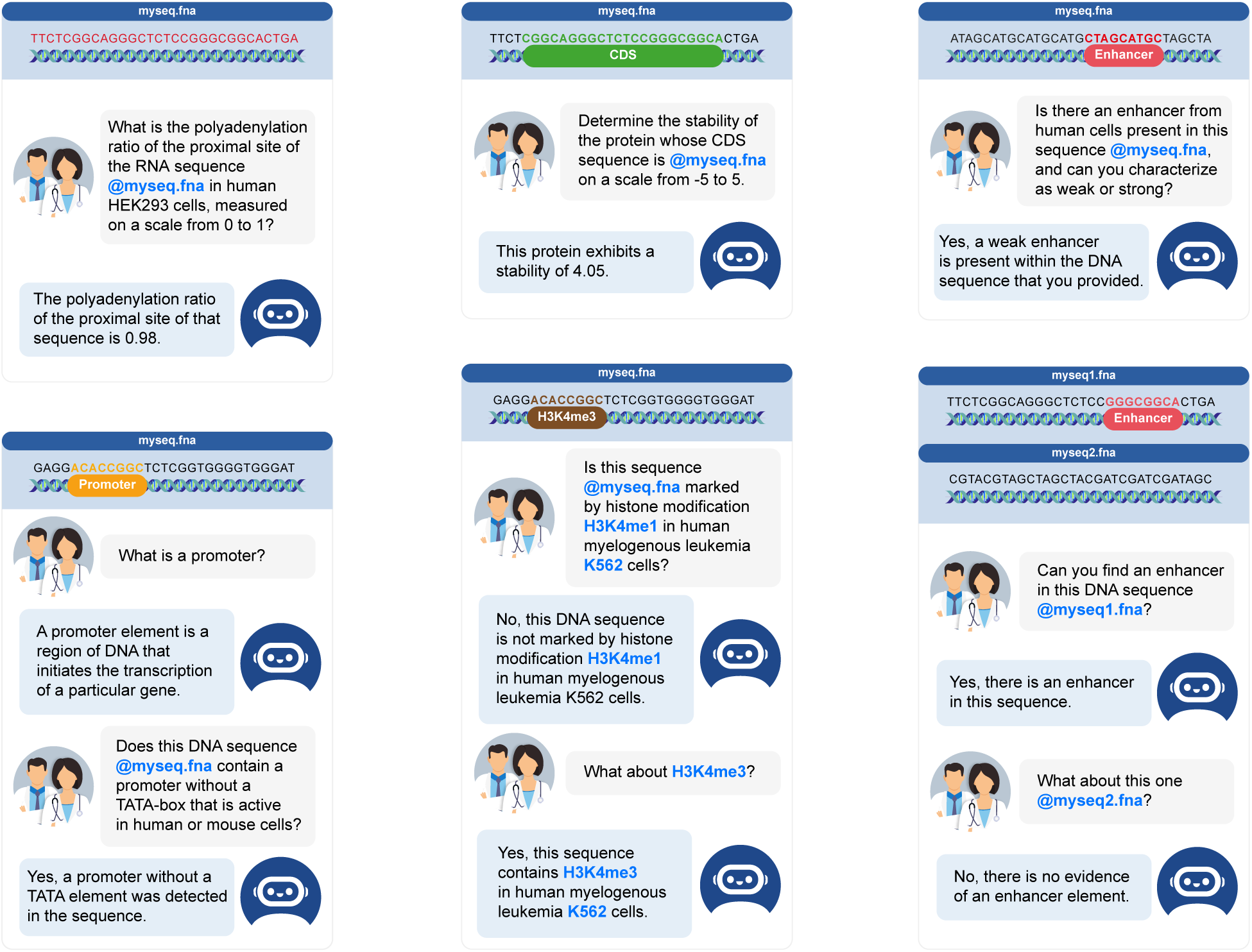
Examples of ChatNT conversations on DNA, RNA and Protein tasks. For each conversation we show the question from the user (white) and the answer of the agent (blue). The projected embeddings of the input DNA sequences are incorporated in the question at the position of *@myseq.fna*.

ChatNT is trained to solve all tasks simultaneously, with a uniform sampling over tasks per batch. Multi-tasking is achieved by ChatNT by prompting in natural language, where the question asked by the user will guide the agent towards the task of interest. Given a text prompt and one or multiple DNA sequences as input, ChatNT is trained to minimize a unified objective for all tasks, which takes the form of the cross-entropy loss between ChatNT predictions and the target answer tokens, as in other instruction-finetuning works [50–52]. This single objective allows to learn seamlessly across tasks without introducing conflicting gradients or scale issues coming from different objectives and loss functions (e.g. Cross-Entropy for classification versus Mean Squared Error for regression). In addition, it allows us to extend the model with additional tasks in the future without requiring changes in the model architecture or training it from scratch. In summary, ChatNT provides a general genomics AI system that solves multiple tasks in a conversational manner, thus providing a new paradigm for genomics models.

In addition to seamlessly integrating multiple types of labeled and experimental data into a single general foundation model, ChatNT is designed to be conversational to enable users to easily interact with it and to use it without requiring a programming background (see examples in Fig. 2). We rely on a frozen English language model, Vicuna 7B [50], that has been instruction fine-tuned from LLaMA [47], ChatNT keeps all the intrinsic conversational capabilities of the language model. Interestingly, we observed that as the training dataset used to build LLaMA already contained a large set of life sciences papers, our agent is also capable to answer multiple questions about genomics such as defining regulatory elements like promoters and enhancers, zero shot i.e. without any additional training data. Additionally, ChatNT can answer numerous non-biology related questions and solve tasks such as summarizing or writing simple programming code. As our approach is general and builds on top of any pre-trained English language model, ChatNT capabilities can improve organically with new and more powerful open-sourced language models. While the conversational capability is an important aspect of ChatNT but is already provided by the respective language model, we focused in this work on demonstrating that the conversational agent ChatNT can solve a wide range of advanced genomics tasks in English with high accuracy.

### ChatNT is the new state-of-the-art on the Nucleotide Transformer benchmark

In order to develop ChatNT and optimize its architecture we created an instructions version of the Nucleotide Transformer benchmark [26] (Supplementary Table 1 and Methods). This collection of genomic datasets is suitable for fast iteration during model experimentation as it contains a varied panel of small-sized datasets and has been extensively evaluated in multiple studies of DNA foundation models [26, 29]. We trained ChatNT to solve all 18 tasks at once and in English and evaluated its performance on test set DNA sequences and questions.

We first used this benchmark to systematically compare the performance of ChatNT with two different projection architectures. The classical way of aggregating information from the encoder in previous multimodal models is to use a trainable projection to convert the encoder embeddings into language embedding tokens, which have the same dimensionality of the word embedding space in the language model [40–42, 49]. In ChatNT we used the Perceiver resampler from Flamingo [41] based on gated cross-attention as projection layer (Supplementary Fig. 1a). Using this projection layer and finetuning both the DNA encoder and the projection on all 18 tasks, ChatNT obtained a new state-of-the-art accuracy on this benchmark with an average Matthew’s correlation coefficient (MCC) of 0.71, 2 points above the previous state-of-the-art Nucleotide Transformer v2 (500M) model (Fig. 3a, Supplementary Fig. 2).

**Figure 3.**
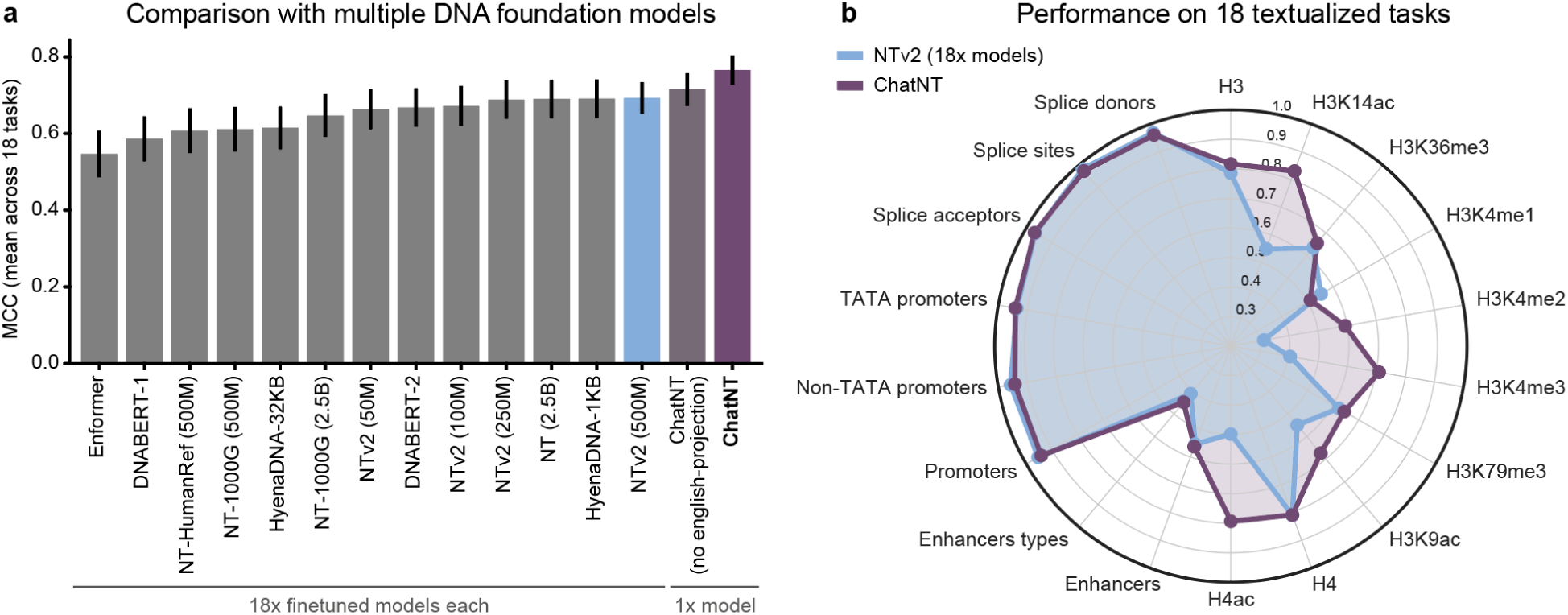
ChatNT achieves a new state-of-the-art accuracy in the Nucleotide Transformer benchmark. **a)** Average performance of ChatNT, ChatNT with no english-aware projection and 13 different genomics foundation models across all 18 tasks of the Nucleotide Transformer benchmark [26]. Bar-plots display the mean MCC over all tasks and the standard error of the mean. **b)** Radar plot depicting the performance of ChatNT in each of the 18 tasks compared with specialized NTv2 models fine-tuned individually on each task.

However, similar to all other projection layers [40, 49, 53], the current implementation of the Perceiver resampler generates the same fixed set of embeddings for the encoder tokens independently of the question asked, and therefore it needs to capture in this set of embeddings all relevant information for every downstream task. We hypothesised that this feature can create an information bottleneck in genomics when scaling the model for multiple downstream tasks given the diversity of potential sequences, from different lengths and species, and biological properties. Therefore, we developed an English-aware Perceiver projection that extracts representations from the input sequence dependent on the English question asked by the user, which allows to leverage contextual information encoded in the input DNA sequences that are relevant for the specific question (Supplementary Fig. 1b; see Methods). We observed significantly improved performance by accounting for the question when projecting the DNA embeddings into the English decoder space (average MCC of 0.77 vs 0.71; Supplementary Fig. 1c,d). This can be explained by the very context- and task-specific information in DNA sequences that we must retain in order to tackle diverse genomics tasks. Since the decoder remains frozen, the projection layer not only needs to bring the sequence embeddings into the embedding space of the English decoder, but also to perform the operations to extract the relevant information from the embedding to answer the question. Our results show that making the projection aware of the question facilitates both aspects thus achieving a better performance and transfer across tasks.

In summary, ChatNT with an English-aware projection (from now on just called ChatNT) achieves a new state-of-the-art accuracy on this benchmark (average MCC of 0.77) in addition to solving all 18 tasks at once (Fig. 3a). Strikingly, ChatNT improves the average performance by 8 points over the previous state-of-the-art Nucleotide Transformer v2 (500M) model, which was used as the DNA encoder within ChatNT (average MCC of 0.77 vs 0.69; Fig. 3a,b). Our results demonstrate that a single unified objective formulated in natural language triggers transfer learning between multiple downstream tasks and helps deliver improved performance.

### A new curated genomics instructions dataset of biologically relevant tasks

Although the Nucleotide Transformer benchmark [26] was very suitable for model experimentation and to debug the system, it misses many tasks of great biological relevance in genomics related to more complex biological processes as well as more recent experimental techniques and tasks that involve quantitative predictions. Therefore we curated a second genomics instructions dataset containing 27 genomics tasks framed in English derived from different studies that cover several regulatory processes (Supplementary Table 2 and Methods). These include tasks related to DNA (21 tasks), RNA (3) and protein sequences (3) from multiple species framed as both binary/multi-label classification and regression tasks. The final instructions dataset contains a total of 605 million DNA tokens, i.e. 3.6 billion base pairs, and 273 million English tokens (including an average of 1,000 question/answer pairs per task) (Figure 1b).

This collection includes a non-redundant subset of tasks from the Nucleotide Transformer [26] and the BEND [54] benchmarks, complemented with relevant tasks from the plant AgroNT benchmark [55] and human ChromTransfer [56]. These benchmarks have been extensively used in the literature, come from different research groups, and represent diverse DNA processes and species. These selected tasks include binary and multi-label classification tasks covering biological processes related to histone and chromatin features, promoter and enhancer regulatory elements, and splicing sites.

We further added state-of-the-art and challenging regression tasks related to promoter activity [55], enhancer activity [11], RNA polyadenylation [20, 21] and degradation [19], and multiple protein properties [57]. These are reference datasets in the respective fields and related to very complex properties of biological DNA, RNA and protein sequences. All RNA and protein tasks are predicted from the corresponding DNA and CDS sequences instead of the RNA and protein sequences, respectively. Getting the matching DNA sequence is trivial for RNA sequences but more challenging for protein sequences due to the complexity of codon usage. Therefore, we used the CDS annotations for protein tasks curated at Boshar et al. [57].

See Figure 2 and 4 for examples of questions and answers for different types of genomics tasks used in our dataset (see also Supplementary Fig. 3, 4, 5). For instance, a training example for an enhancer classification task would be “*User: Is there an enhancer from human cells present in this sequence @myseq.fna, and can you characterize as weak or strong? ChatNT: Yes, a weak enhancer is present within the DNA sequence that you provided.*”, where the projected embeddings of the candidate DNA sequence are inserted at the @myseq.fna position. Regression tasks are also framed in English and the agent needs to write the digits corresponding to the requested quantity: for example “*User: Determine the degradation rate of the mouse RNA sequence @myseq.fna on a scale from -5 to 5. ChatNT: The measured degradation rate for this sequence is 2.4.*” (see Methods for details on the quantitative scale). The loss is equally computed as the cross-entropy loss between the predicted and the target answer tokens. For performance evaluation, we extract the digits from each answer and test their correlation with the ground-truth values.

**Figure 4.**
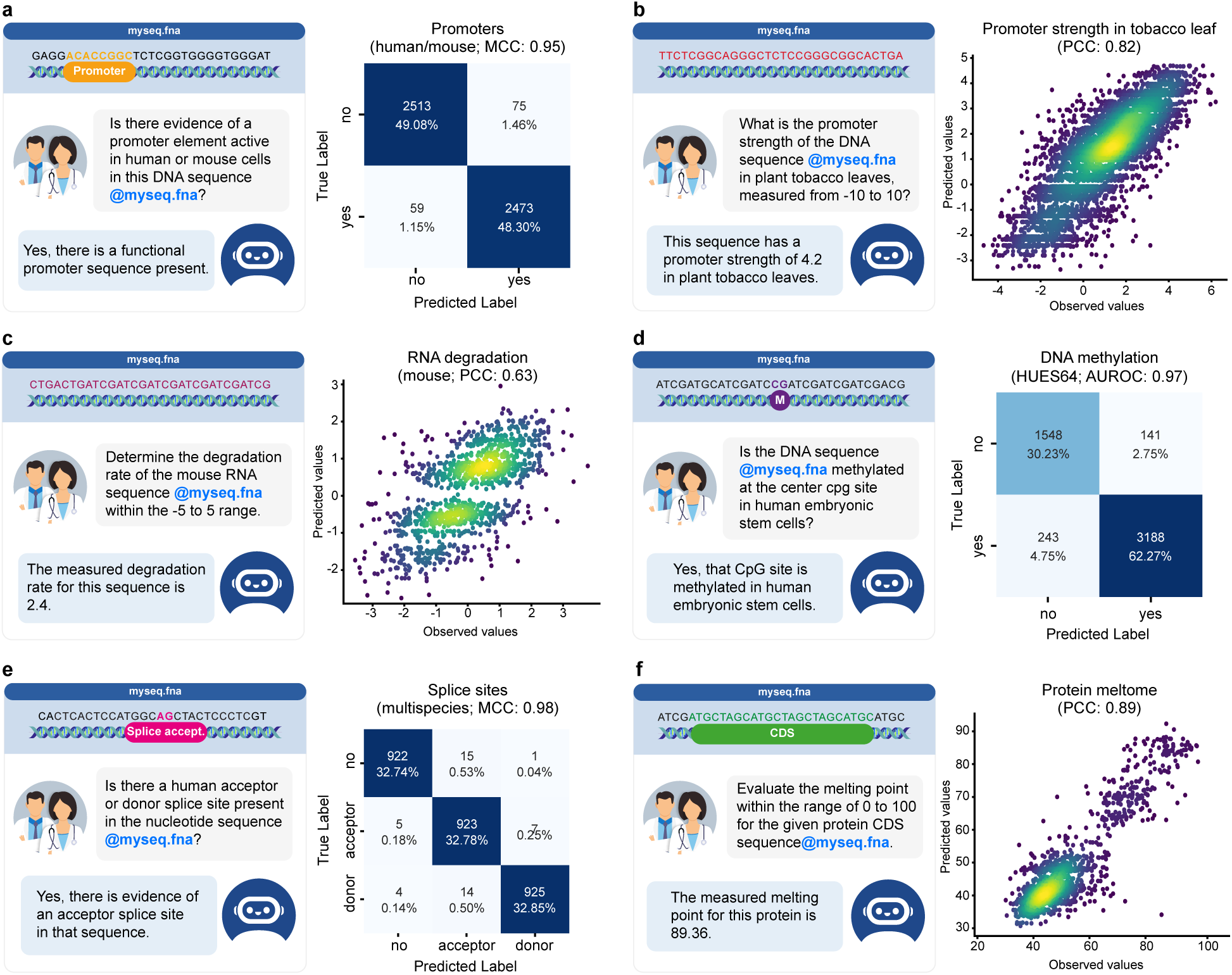
Examples of prediction performance and conversations for a subset of genomics, transcriptomics, and proteomics tasks. **a, d, e)** Left: example of conversation for the respective binary or multi-label classification task. Right: heatmap displaying the confusion matrix comparing the predicted labels of ChatNT and observed labels. The performance metric is reported. **b, c, f)** Left: example of conversation for the respective regression task. Right: scatter-plot comparing the predictions of ChatNT and observed values. Pearson correlation coefficient (PCC) is reported.

In summary, this curated set of tasks provides a general perspective of the capabilities and usefulness of our model in different biological sequence domains. We train ChatNT as a general agent to solve all 27 genomics tasks at once and in English, and compare its performance with the state-of-the-art specialized model for each task (see Methods).

### ChatNT achieves high performance on multiple tasks across different genomics processes and species

We first evaluated the performance of ChatNT on the 21 tasks related to different DNA processes from yeast, plants, fly, mouse, and human. ChatNT is competitive with the performance of the different specialized models that were fine-tuned directly on each of these individual tasks (Fig. 4a,b,d,e and 5a,c). In particular, we obtained an improved performance on the detection of human enhancer types. Still, we observed significantly reduced performance for enhancers from plant species when compared with the state-of-the-art AgroNT model fine-tuned specifically on this task [55]. Since AgroNT was pre-trained on genomes from 48 diverse plant species, improving the encoder used in ChatNT might lead to improved performance on this type of tasks.

**Figure 5.**
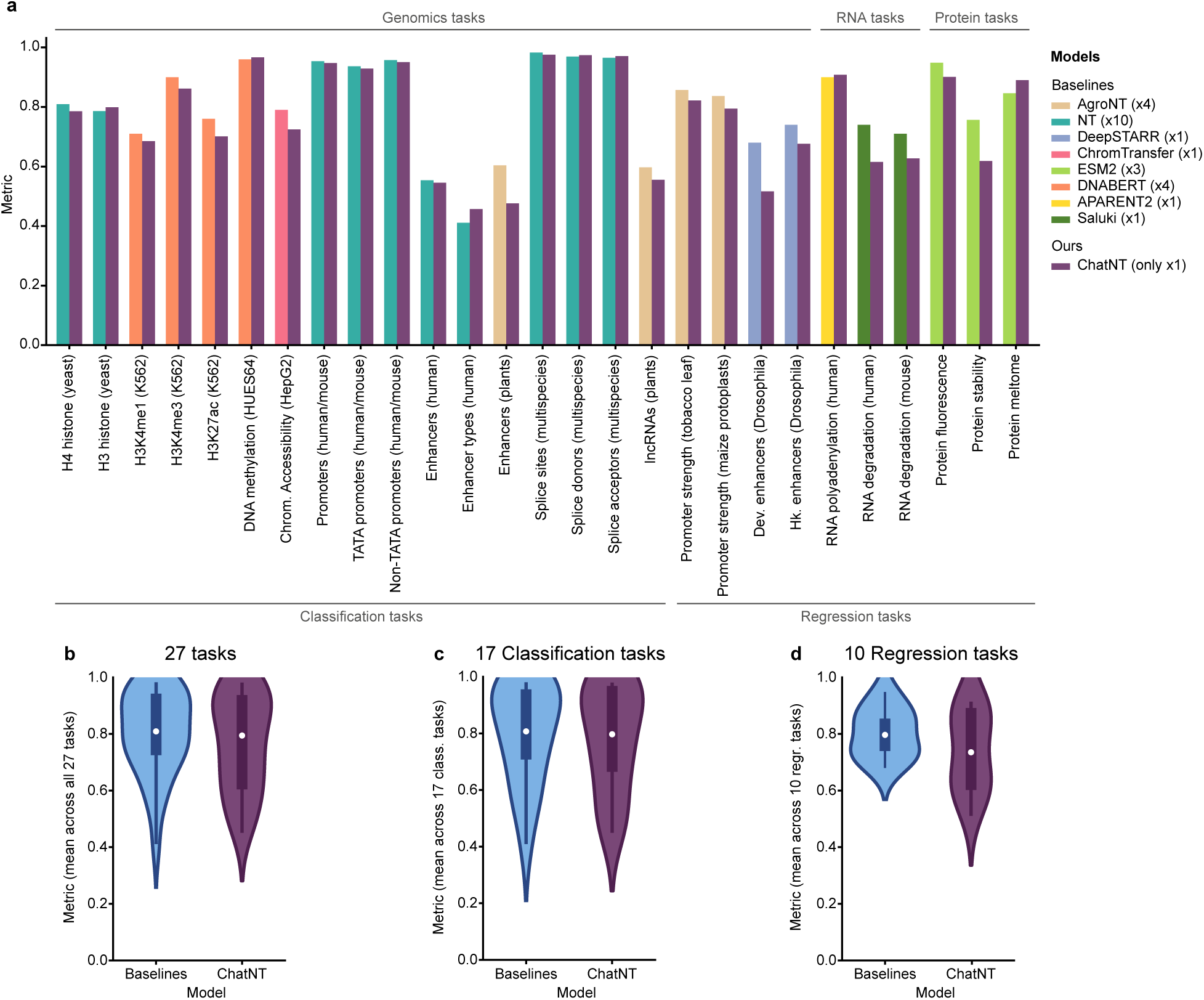
Comparison of ChatNT vs. domain specific baselines over genomics, transcriptomics and proteomics tasks. **a)** Barplots with the performance of ChatNT compared with respective baselines per task. The metric used for each task is the same used in the respective baseline study (Supplementary Table 2). **b-d)** Comparison between ChatNT and baselines for (**b**) all tasks, (**c**) classification tasks and (**d**) regression tasks. Metrics are the same as in (**a**). The box plots mark the median, upper and lower quartiles and 1.5× interquartile range (whiskers); outliers are shown individually.

As ChatNT solves the tasks in English, it can seamlessly handle binary and multi-label classification tasks. By extracting the term predicted by ChatNT in the answer, we can quantify its predictive performance. As we show for some examples in Fig. 4, ChatNT accurately identifies input sequences with human or mouse promoters (Fig. 4a), with CpG sites methylated in human embryonic stem cells (HUES64 cell line; Fig. 4d), and with splice acceptor and donor sites (Fig. 4e).

ChatNT is also able to solve quantitative tasks by writing the digits of the predicted score. We observed competitive performance on predicting promoter activity in plants, namely tobacco leaves (Fig. 4b) and maize protoplasts, but significantly reduced performance on *Drosophila* enhancer activity over the state-of-the-art DeepSTARR model [11] (Fig. 5a). Importantly, the distributions of the predicted digits correlate well with the original scores (Fig. 4b). This capability to proficiently address regression tasks is of paramount importance in biology, and is particularly significant in light of the acknowledged limitations and unreliability of numerical processing in language models [58, 59]. Still, we observed a reduced average performance on regression tasks over classification ones, likely due to the difference in complexity and classification tasks being more represented in the training set. We assume that this might be solved by improving the balance between classification and regression tasks during training, through either a weight loss or a task sampling frequencies curriculum [60].

### ChatNT solves transcriptomics and proteomics tasks

ChatNT is built with a flexible architecture that allows it to handle any type of biological sequence that can be processed with our DNA encoder, the Nucleotide Transformer [26]. To showcase its generalization, we have included in the new genomics instructions dataset three RNA and three protein regression tasks (Supplementary Fig. 4, 5). These include predicting RNA polyadenylation and degradation rates as well as different protein features. Examples of conversations used for model training are: “*User: What is the measured polyadenylation ratio of the proximal site of the RNA sequence @myseq.fna in human HEK293 cells, considering a range from 0 to 1? ChatNT: That sequence has a polyadenylation ratio of the proximal site of 0.69.*” and “*User: Specify the melting point of the protein with the given coding sequence (CDS) @myseq.fna within the 0 to 100 range. ChatNT: This protein demonstrates a melting point of 80.81.*”. The performance of ChatNT was compared to the state-of-the-art specialized models APARENT2 for polyadenylation [21], Saluki for RNA degradation [19], and ESM2 for the protein tasks [48] (Supplementary Table 2).

Overall, we observed good performance for ChatNT on the test sets of the 6 RNA and protein tasks, with Pearson correlation coefficients (PCCs) between 0.62 and 0.91 (Fig. 4c,f, 5a). ChatNT outperformed the specialized models for the prediction of proximal polyadenylation site ratio (PCC of 0.91 vs 0.90) and protein melting points (PCC of 0.89 vs 0.85). Regarding the RNA degradation tasks in human and mouse, ChatNT obtained a PCC of 0.62 and 0.63, ten points below the specialized Saluki model [19] (PCC of 0.74 and 0.71). ChatNT also obtained competitive performance with the state-of-the-art protein language model ESM2 [48] on the two other protein tasks related to protein fluorescence and stability. Although ChatNT cannot yet outperform every specialized model on RNA and protein tasks, we show that it can already handle such tasks and achieve high performance using the DNA foundation model Nucleotide Transformer as a DNA encoder. ChatNT’s flexible architecture allows to plug-in different encoders, such as language models specialized for RNA [61–64] and protein domains [48], which should reduce the gap to specialized deep learning models in the transcriptomics and proteomics fields and improve the capabilities and generalization of ChatNT towards a unified model of biology.

### Assessing the confidence of ChatNT answers

ChatNT is built to assist and augment scientists and researchers in their daily research. As such, its performance and reliability are paramount. However, in contrast to standard machine learning models that return probabilities or quantitative scores, ChatNT directly answers questions, preventing the user to get a sense of its confidence and thus reducing its practical value for sensitive applications. This is an important challenge and common to all current conversational agents [38–40]. To address this, we introduce a novel way to assess the confidence of our agent for binary classification tasks. Instead of generating directly answers to the binary classification question for a given sequence, we compute the model perplexity for that question over examples of both positive and negative answers. We make sure that these selected answers were not included in the model training dataset. Those perplexity values towards positive and negative answers are then used to derive logits and probabilities for each class for the candidate question. This method allows us to derive probabilities from ChatNT for each question example, similar to standard classifiers, and we refer to it as perplexity-based classifier (Fig. 6a).

**Figure 6.**
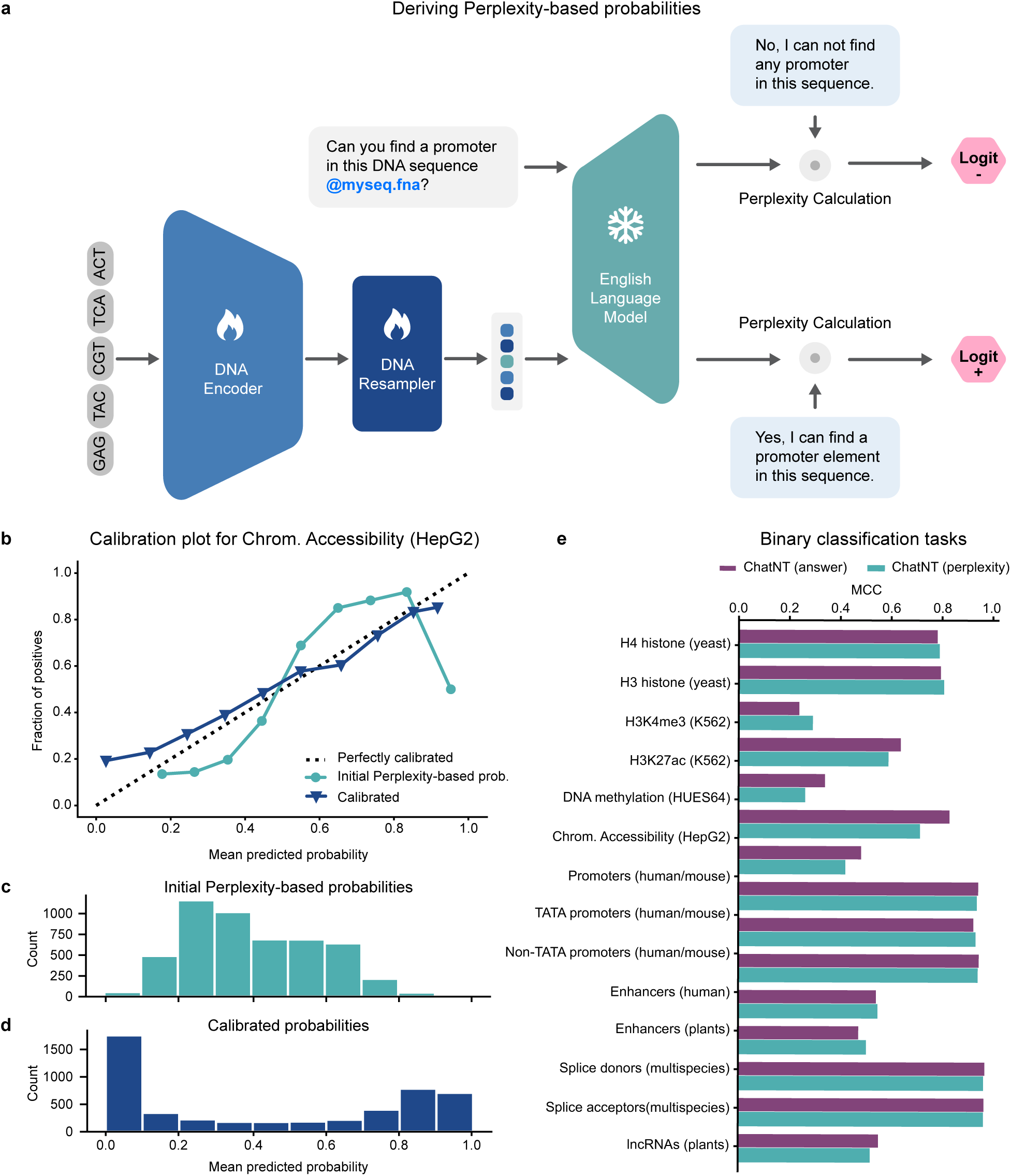
Perplexity-based method to calibrate the confidence of ChatNT answers while preserving performance. **a)** Cartoon describing the perplexity-based classifier based on ChatNT answers. **b)** Calibration plot for the task of human chromatin accessibility (cell line HepG2). Scatter-plot comparing the predicted probability and fraction of positives over ten bins for the original (orange) and calibrated (purple) perplexity-base classifiers. **c)** Histogram with the predicted probability over ten bins for the original perplexity-base classifiers. **d)** Histogram with the predicted probability over ten bins for the calibrated perplexity-base classifiers. **e)** Comparison of the performance (MCC) of the ChatNT answers (yes vs no) and its derived perplexity-based probabilities for all binary classification tasks.

Computing probabilities enables us to assess the calibration of the model, i.e. the correlation between the predicted probability, its confidence, and the accuracy of its prediction. We say that a model is well calibrated when a prediction of a class with confidence p is correct 100p % of the time. We computed the ChatNT perplexity-based probabilities for all binary classification tasks. In Figure 6b-d we show an example of a calibration plot based on the predictions for the chromatin accessibility task. We observe that our model is well calibrated for low- and high-confidence areas, but less in medium-confidence ones. For instance, examples predicted with a probability of 0.9 are correctly predicted 90% of the time while examples predicted with probability 0.5 are correctly predicted only 25% of the time. To improve this, we show that we can calibrate our model by fitting on the training set a Platt’s model [65], to improve the confidence of the model across all ranges of predictions (Fig. 6b-d). This calibration step is performed for all binary classification tasks. Overall, we achieve the same performance for ChatNT across tasks using these perplexity-based predictions (Fig. 6e) but with improved calibration. As a consequence, our approach can accurately measure the predictive performance of a language model in addition to effectively assessing its uncertainty level. This technique, while being general, should also be beneficial to other language model fields.

## Discussion

We presented ChatNT, the first multimodal conversational agent that can handle DNA, RNA and protein sequences and solve multiple biologically relevant downstream tasks. We built and curated the first datasets of genomics instructions tasks including binary and multi-labels classification and regression tasks spanning different species and genomics processes. Tasks relative to transcriptomic and proteomic processes were also included to demonstrate the versatility and generality of this approach across domains. ChatNT achieves a new state-of-the-art on the Nucleotide Transformer benchmark [26] and demonstrates a performance on par with specialized models on our new set of 27 tasks. Importantly, unlike conventional approaches requiring a specialized model for each task, ChatNT solves all tasks within a unified model in addition to offering a simple and natural chatbot interface for people to use the model. We also introduced a new technique to probe the confidence of language models for binary classification tasks and used it to calibrate them when needed. Altogether, ChatNT is the first proof that natural language LLMs can be extended to process bio-sequence modalities, displaying not only conversational capabilities but also answering accurately multiple biologically relevant questions.

To extract the complex information from DNA sequences that is needed to solve all tasks in a single unified model, we introduced a novel architecture based on the Perceiver resampler [41] to resample and project DNA embeddings into the natural language embedding space. We identified an information bottleneck issue that arises from the diversity of tasks, species and biological processes encoded in DNA sequences, and we showed how to solve it by conditioning the projection on the question asked. This conditioning allows the projection module to extract from the DNA embeddings the right amount of information to solve the task at hand, as we show by the improved performance over a projection module that is not conditioned on the question.

In this work, we decided to focus on situations where a user, such as a researcher or scientist, is interested in detecting molecular phenotypes or computing quantitative properties for a given DNA sequence. While we believe this encompasses an already significant number of practical use-cases, it would be interesting to expand the agent capabilities to handle other typical bioinformatics pipelines. Such pipelines could include calling tools to compute statistics about the sequences, aligning the sequences to a reference database to compute multiple sequence alignments, query external databases for additional information about the sequences, or to recursively call the ChatNT model over a FASTA file containing multiple sequences and generating a summarized table results with its corresponding analysis. This is supported by the success of external tools in large language models such as Toolformer [66], LLaVA-Plus [49], geneGPT [67] or GPT-4 [39]. Such pipelines could also benefit from ChatNT’s capability to handle several sequences at the same time in order to reduce the inference compute cost. Replacing ChatNT’s current English decoder by larger models and/or models fine-tuned using Reinforcement Learning Human Feedback (RLHF) such as Llama2-chat 70B [52] could also help extending the model capabilities in these directions as well as improving its overall usefulness.

The capabilities of ChatNT have been demonstrated for DNA sequences using a pre-trained DNA foundation model, the Nucleotide Transformer [26]. As shown in our experiments, working with DNA sequences allows to tackle tasks not only in genomics but also transcriptomics and proteomics, the latter using the corresponding CDS region. However, our approach could be easily extended to integrate encoders from other omics modalities such as RNA [61–64] and protein [48, 68] language models to work natively with RNA and aminoacid sequences. Through our positional tag system that supports multiple sequences, one could simply add an arbitrary number of encoders and train their respective projections to combine different omics and modalities within the same questions. We envision that such approach could expand even further the capabilities and performance of our model by achieving superior transfer learning across modalities.

This work serves as the first proof-of-concept that it is possible to build multimodal bio-sequence/ English conversational agents that can solve advanced, biologically relevant tasks, and is meant to lay a first set of foundations to build future highly-capable agents that understand biological sequences and principles. Similar to the developments in NLP [52, 69–71] and multimodal models [72], we expect new capabilities such as zero-shot performance to emerge through developments on two main fronts: (1) scaling the number of tasks by including examples from diverse biological processes, tissues, individuals and species [73, 74]; and (2) integrating more data modalities, such as RNA and protein sequences, imaging data and health records from individuals. When such capabilities emerge, it will be of the highest importance to carefully assess model safety and robustness, for instance through red teaming [75]. As such, ChatNT represents an important step along the trajectory towards general purpose AI for biology and medicine [76].

## Methods

### ChatNT model

#### Architecture

The ChatNT is a multimodal agent that takes as input one or multiple DNA sequences and an English prompt and returns a distribution over English words that is used to auto-regressively produce an answer in English. We introduce a DNA English token placeholder <DNA> that is added in the input English prompt for the user to refer to the DNA sequence. The architecture is also extended to handle several DNA sequences. In this case, each DNA sequence is processed independently by the DNA encoder and the input English prompt is expected to contain as many DNA English token placeholders as sequences are inputted.

The ChatNT architecture is made of three parts: a pre-trained DNA encoder, a projection model that projects the DNA embeddings into the English tokens embedding spaces and a pre-trained English decoder. While our architecture is general and could work with any choice of DNA Encoder and English decoder, we decided to use the pre-trained Nucleotide Transformer v2 (500M parameters) [26] and Vicuna-7b (instruction fine-tuned Llama model with 7B parameters) [50] models, respectively. During training, we keep the English decoder frozen and update only the weights of the DNA encoder and the projection model. The projection model is initialized from scratch at the beginning of the training.

The DNA Encoder processes the DNA sequence and returns one embedding vector per input token, one token representing a nucleotide 6-mers in the case of the Nucleotide Transformer model. We note L the number of nucleotides in the DNA sequence and N the number of DNA tokens (with roughly 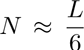). Every in-6 put DNA sequence was padded if needed until a final length of 2, 048 tokens, representing approximately 12*kb*. As the output embedding dimension of the DNA encoder can be different from the words embedding dimensions of the English language model we first use a dense neural network to project each DNA token embedding to the English word dimension. In a second phase, we use a Perceiver resampler architecture [41] that uses cross-attention between the projected DNA tokens embeddings and learnable queries, to re-sample the *N* DNA tokens embedding to *K* embedding vectors (Supplementary Fig. 1a). We have adapted this Perceiver resampler to include an additional cross-attention step between the learnable queries and the English question in order to extract context-dependent representations from the DNA sequence (Supplementary Fig. 1b).

On the other hand, the English prompt is tokenized and English tokens embeddings are produced for each tokens. The *K* resampled DNA embedding vectors are then inserted in place of the DNA sequence placeholder tokens in the English input sequence. In the case of multiple input DNA sequences, these operations are applied consecutively and independently for each DNA sequence. We experimented with several values of *K* in practice and we observed that low values such as 1 or 4 are not enough for the DNA encoder to impact the behavior of the frozen English decoder. We found *K* = 64 to provide a good trade-off between the input length of the English decoder and the performance in practice.

During inference, the DNA encoder embeddings for the DNA sequences are computed only once. The inference is done autoregressively by predicting sequentially each new token until an end of sequence token is predicted. The key, queries and values of the English decoder are cached during generation to avoid computing unnecessary operations. We use temperature sampling with a temperature of *τ* = 0.001.

The whole codebase of the ChatNT has been developed in Jax [77] using Haiku [78] for neural networks implementation. All trainings were performed on a cluster of 8 GPU H100 instances and evaluations of the model can be done in a single GPU A100-80gb. All trained parameters from the DNA encoder and perceiver projection as well as optimizer accumulators and all frozen parameters from the English decoder are stored and updated in float32.

#### Training

ChatNT was trained using Adam optimizer [79] with *lr* = 3*e^−^*^5^ and default settings for other hyperparameters: β_1_ = 0.9, β_2_ = 0.999, ε = 1*e^−^*^8^, ε_root_ = 0.0. We used gradient clipping of 1 and accumulated gradients over a batch size of 65, 536 tokens, equivalent to 256 samples. We used an uniform sampling over tasks per batch such that each batch has the same proportion of samples per task. We trained the model on the 27-task dataset for 2B tokens (7.8M samples) on a cluster of 8 GPU H100 over 4 days.

#### Hyperparameters

Below we describe all hyperparameters for the different parts of ChatNT.

|  | DNA encoder | Perceiver Resampler | English decoder |
| --- | --- | --- | --- |
| Number of layers | 29 | 3 | 32 |
| Number of heads | 16 | 20 | 32 |
| Embedding dimension | 1024 | 4096 | 4096 |
| Feed forward dimension | 4096 | 11008 | 11008 |
| Activation type | swish | GeLu | swiGlu |
| Positional encoding type | RoPe | RoPe | RoPe |
| Total number of parameters | 500M | 800M | 7B |
| Input tensor shape | (1, 2048) | (1, 2048, 1024) | (1, 1024, 4096) |
| Output tensor shape | (1, 2048, 1024) | (1, 64, 4096) | (1, 1024, 32000) |
| Float Precision | Float32 | Float32 | Float32 |
| Initialization | Pre-trained (NT-v2-500m) | From scratch | Pre-trained (vicuna-7b) |
| Update | Updated | Updated | Frozen |

#### Evaluation

Evaluating the performance of ChatNT can be done in a single GPU A100 in batches of 32 samples and takes 1:40 minutes to generate a maximum of 40 tokens per sample (13 tokens per second). For each task, we evaluated ChatNT on upmost 5, 000 sampled test samples and report the metric used in the respective benchmark study (Supplementary Table 1 and 2).

### Genomics instructions datasets

#### Instructions for the Nucleotide Transformer benchmark

We created an instructions version of the Nucleotide Transformer benchmark [26] (Supplementary Table 1). To convert the DNA sequence datasets into instructions datasets, we curated dozens of English questions and answers for each task and sampled a question/answer pair per input DNA sequence. We used the DNA token placeholder <DNA> in the question when referring to the input DNA sequences. The answer contains the classification label for the respective input sequence. We converted all 18 binary/multi-label classification datasets into diverse question/answer instructions for each DNA sequence. We provide for each task train and test sets containing different DNA sequences as well as different questions to assess the performance and English generalization capabilities of the model. We kept the same train and test sets as the original dataset.

#### New curated genomics instructions dataset of biologically relevant tasks

The new genomics instructions dataset created here contains a set of 27 tasks framed in English derived from different studies (more details in Supplementary Table 2). It covers several regulatory processes related to DNA (21 tasks), RNA (3) and protein sequences (3). These tasks are derived from multiple species, including human, mouse, fly and plants. Among all tasks there are 15 binary classification, 2 multi-label classification and 10 regression tasks. The number of training examples per task ranges from 5.5*K* to 3*M*. See Supplementary Information for all details on the data references and processing for each specific task.

We converted the DNA sequence datasets into instructions datasets as described above for the Nucleotide Transformer benchmark. The answer contains the classification label or regression score (up to decimal cases) for the respective input sequence. In addition to simple examples with a single turn of question/answer with a single sequence, we also added more complex examples with multiple turns with consecutive questions that can be related or not, and exchanges where the question refers to multiple sequences. The final genomics instructions dataset contains a total of 605 million DNA tokens, i.e. 3.6 billion base pairs, and 273 million English tokens (including questions and answers).

We obtain for each task train and test sets containing different DNA sequences as well as different questions to assess the performance and English generalization capabilities of the model.

#### Baselines for the genomics tasks

For each of the 27 genomics tasks, we compared the performance of ChatNT with the state-of-the-art method for the respective dataset. These included the convolutional neural networks DeepSTARR [11], ChromTransfer [56], APARENT2 [21] and Saluki [19]; and the fine-tuned foundation models based on Nucleotide Transformer [26], agroNT [55], DNABERT [27] and ESM2 [48]. We used different performance metrics per task to follow the same metric used in the respective studies. Details on the baseline method and performance metric per task can be found in Supplementary Table 2. Most baseline performance metrics were directly retrieved from the respective papers. Only for ESM2 we had to rerun them on the updated dataset versions.

### Calibration of ChatNT predictions

We developed an approach to assess and calibrate the confidence of ChatNT answers for binary classification tasks.

For a given binary classification task, we select *N* examples of positive and negative answers each, selected from the respective task’s test set. We note these examples respectively 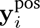 and 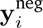 where 0 *≥ i* > *N*. Then, *i i* for a given question **x** and DNA sequence **s**, we compute the average perplexity of the model over the positive and negative examples respectively. We denote these two values as 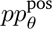(**x**, **s**) and 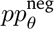(**x**, **s**), respectively, where θ represents the ChatNT weights tensor. We compute them as follow:

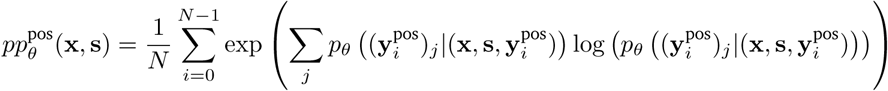

where 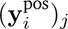 denotes the j-th token of answer 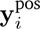 and 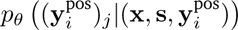 returns the probability of token *j* given the question, DNA sequence and tokens from the answers up to the j-th one according to ChatNT. The negative perplexity values are computed similarly over negative answers.

Those perplexity values towards positive and negative answers represent a measure of how well the model aligns the question to those answers. We inerpret them directly as logits and use a softmax transformation to compute probabilities for the respective class for the input question. This method allows to derive probabilities from ChatNT for each question example. We applied this approach to 1, 000 test examples per task.

To calibrate those predictions, we first compute perplexity-based probabilities to 10, 000 training examples as our calibration dataset and use them to fit a Platt’s model [65]. More specifically, we use logistic regression from scikit-learn [80] as the calibrator model and trained it with the following parameters with an inverse regularization factor *C* = 0.1 and with the lbfgs solver. The logistic regression model learns to map the perplexity-based probabilities from ChatNT onto a more accurate scale. We then apply this model to calibrate the probabilities of the 1, 000 test examples mentioned above.

As metrics, we computed both Area under the ROC Curve (AUROC) and MCCs for both the original perplexity-based probabilities and the calibrated ones.

## Data availability

All input data are freely available from public sources referenced in the respective Methods section. All genomics instructions datasets prepared for training ChatNT are available as supplementary files, including the DNA sequences, questions and answers of each dataset. We also provide all questions and ChatNT answers on the test set sequences used to evaluate its performance on the different tasks.

## Code availability

We provide the pseudocode of the algorithmic steps and key concepts underlying our multi-modal model in the section Supplementary Pseudocode.

## Acknowledgments

We thank Aliou Kayantao for his help improving the figure panels and Boulbeba Mallouli and his team for help on the ChatNT interface. We thank Evan Trop, Javier Mendoza-Revilla, Maša Roller, Nicolas Lopez Carranza, Liviu Copoi and Marcin Skwark for helpful discussions. We would also like to thank Lida Rosseló for help on the project management side of this research project.

## Competing interests

G.R., B.P.d.A., H.D-T., M.L., A.L., K.B. and T.P. are employees of InstaDeep LTD. C.B., L.H., P.P., M.L. and U.S. are employees of BioNTech LTD.

## Supplementary Information

### Genomics instructions dataset

#### Histone modifications

As representatives for yeast histone datasets, we used as tasks the presence of H3 and H4 histones along the yeast genome derived from Chip-Chip experiments [81]. The processed data for the yeast H3 and H4 tasks was retrieved from the Nucleotide Transformer benchmark [26]. We used MCC as performance metric per histone type.

As representatives for human histone datasets, we used the abundance of the histone modifications H3K4me1, H3K4me3 and H3K27ac along the human genome in the model cell line K562. Training and test DNA sequences and respective positive and negative labels were obtained from the BEND benchmark study [54]. Each input sequence is of length 512bp and is assigned a positive label if a histone bound to it carries the respective mark. We reduced the size of the dataset for practical reasons by downsampling the negative sequences to twice the number of positive sequences. We used AUROC as performance metric per histone modification.

#### Chromatin accessibility

We retrieved an example of a chromatin accessibility prediction task from ChromTransfer [56], selecting data from the cell line HepG2 since it was the most challenging task in the dataset. We used their fine-tuning dataset based on ENCODE data with input sequences of 600bp. Positive sequences were defined as regions that were only accessible in that cell line among the six cell lines considered in the study (n=31, 211 for HepG2), while negatives (n=54, 995) were sampled from the positives of the other cell lines and other regulatory regions from ENCODE. We used the F1 score as performance metric.

#### DNA methylation

We collected DNA methylation processed data for the human embryonic cell line HUES64 from the BEND benchmark study [54]. Each input sequence is of length 512bp and contains a CpG site at the center that is either methylated or not. Similarly to histone marks, we reduced the size of the dataset by downsampling the negative sequences to twice the number of positive sequences. We used AUROC as performance metric similar to the BEND benchmark.

#### Human and mouse regulatory elements

We retrieved the dataset of human and mouse promoter sequences used in the Nucleotide Transformer benchmark [26], originally derived from DeePromoter [82]. We considered sequences of 300bp that span 249bp upstream and 50bp downstream of transcription start sites. This resulted in 29, 597 promoter regions, of which 3, 065 contain and 26, 532 do not contain a TATA-box motif. We used the same negative sets, ending up in a total of 59, 194 sequences. We used these sequences for three different binary classification datasets: classifying sequences as promoters (NT_promoter_all), promoters without a TATA-box motif (NT_promoter_no_tata), and promoters with a TATA-box motif (NT_promoter_tata).

For human enhancer prediction tasks we used the enhancer dataset from the Nucleotide Transformer benchmark [26], curated priory at [83]. This dataset contains enhancer (strong or weak) and non-enhancer sequences of 200bp each. We derived two tasks from this dataset: a binary classification task for predicting enhancers (strong and weak combined; NT_enhancers) and a multi-label classification task for classifying a sequence as a strong enhancer, weak enhancer or not an enhancer (NT_enhancer_types). Each dataset contained 14, 968 training sequences and 400 test sequences.

#### Multi-species splice sites

We collected the splice site prediction tasks from the Nucleotide Transformer benchmark [26]. These were based on two original datasets.

We used a dataset originally from SpliceFinder [84] that contains a training set (n=27, 000) of 400bp sequences that contain donor, acceptor, or non-splice sites detected in human genes. The test set (n=3, 000) contains similar types of sequences from human but also additional species: mouse, rat, fly and zebrafish. This dataset was transformed in a multi-label classification task with labels being acceptor, donor or none (NT_splice_sites_all).

We used two additional binary classification tasks for the predictions of donor (NT_splice_sites_donors) or acceptor (NT_splice_sites_acceptors) splice sites. This task was derived from the Spliceator dataset [16], based primarily on the G3PO database, which included sequences from 147 phylogenetically diverse organisms (ranging from protists to primates, including humans). All sequences were 600bp and were labeled as positive if they included a splicing site at the center (i.e. an acceptor or donor site, respectively). The NT_splice_sites_donors dataset contained 19, 775 training and 2, 198 test sequences while the NT_splice_sites_acceptors dataset contained 19, 961 training and 2, 218 test sequences.

#### Plant enhancers

We retrieved the binary classification task for predicting enhancers in the cassava plant (*Manihot esculenta*) seedlings from the AgroNT benchmark [55]. This is a balanced and GC-matched dataset of 1000bp sequences that contain or do not contain enhancers. Sequences from every chromosome except 9 and 17 were used for training (n=16, 852) while sequences from the chromosome 17 were used for testing (n=812).

#### Plant lncRNAs

For the binary classification task of predicting plant long non-coding RNAs (lncRNA), we used the dataset of *Sorghum bicolor* from the AgroNT benchmark [55]. This dataset contains lncRNA sequences with a length smaller than 6, 000bp labelled as positives and length- and GC-matched mRNA sequences labelled as negatives. We used the same training (8, 654) and test (734) sets.

#### Plant promoter strength

The promoter strength dataset from plants was derived from the AgroNT benchmark [55]. This dataset contains 170bp promoter sequences from three different plant species whose strength was tested in tobacco leaves and maize protoplasts. We used the resultant quantitative values for the two different promoter strength regression tasks.

#### Enhancer activity

For tasks related to enhancer activity we considered the DeepSTARR dataset [11]. The dataset is composed of 484, 052 DNA sequences of length 249bp, each measured for their quantitative enhancer activity towards a developmental or a housekeeping promoter in *Drosophila melanogaster* fruitfly S2 cells. We considered these two measures as two regression tasks and used the same training (402, 304) and test (41, 184) set sequences.

#### RNA polyadenylation

We retrieved the data for the RNA polyadenylation task from APARENT2 [21]. This dataset was originally derived from Bogard et al. [20] and we applied the same processing as in APARENT2 to make the training data more uniform. It contains 185bp sequences with randomized proximal polyadenylation signal (PAS) sequences that were tested within 12 diverse 3UTR contexts in an MPRA experiment. The objective is to predict the total isoform proportion of a far-away competing distal PAS. This regression task contains 3.3 million training sequences and 80, 000 sequences testing.

#### RNA degradation

We retrieved the data for the human and mouse RNA degradation tasks from Saluki [19]. This dataset contains processed half-lives for different human and mouse RNA sequences. We used the cross-validation dataset from fold 0 and removed RNA sequences longer than 12kb. This resulted in 10, 377 training and 1, 297 testing human sequences, and 10, 989 training and 1, 374 testing mouse sequences.

#### Protein tasks

We retrieved three different protein tasks related to protein fluorescence, stability and meting point, all predicted from the respective CDS sequence, from Boshar et al. [57].

Protein fluorescence: Estimating the fitness landscape of protein variants which are many mutations away from the wildtype sequence is one of the core challenges of protein design. This task evaluates a model’s ability to predict log-fluorescence of higher-order mutant green fluorescent protein (GFP) sequences. Original data is from an experimental study of the GFP fitness landscape [85]. Inspired from the TAPE and PEER benchmarks [86, 87], we restrict the training set to amino-acid sequences with three or fewer mutations from parent GFP sequences, while the test set is all sequences with four or more mutations.

Protein stability: It is important for models trained on diverse sequences to be able to accurately predict a small region of the fitness landscape. This task evaluates how well models predict stability around a small region of high-fitness sequences. Coding sequences and labels were taken from the supplementary material of the original experimental study [88]. Labels indicate a peptide’s ability to maintain structure at increasing levels of protease, which serves as a proxy for stability.

Protein melting point: Predicting protein melting point can be a challenging task as even single residue mutations can have large impacts [89]. Melting point prediction is a sequence-level regression task that evaluates a model’s ability to predict a measure of melting temperature. We follow the same “mixed” splits described in FLIP [22] which seek to avoid over-emphasis of large clusters. Sequences are clustered at 20% identity with 80% of clusters assigned to the train dataset and 20% of clusters assigned to the test dataset.

## Supplementary Tables

**Supplementary Table 1.**
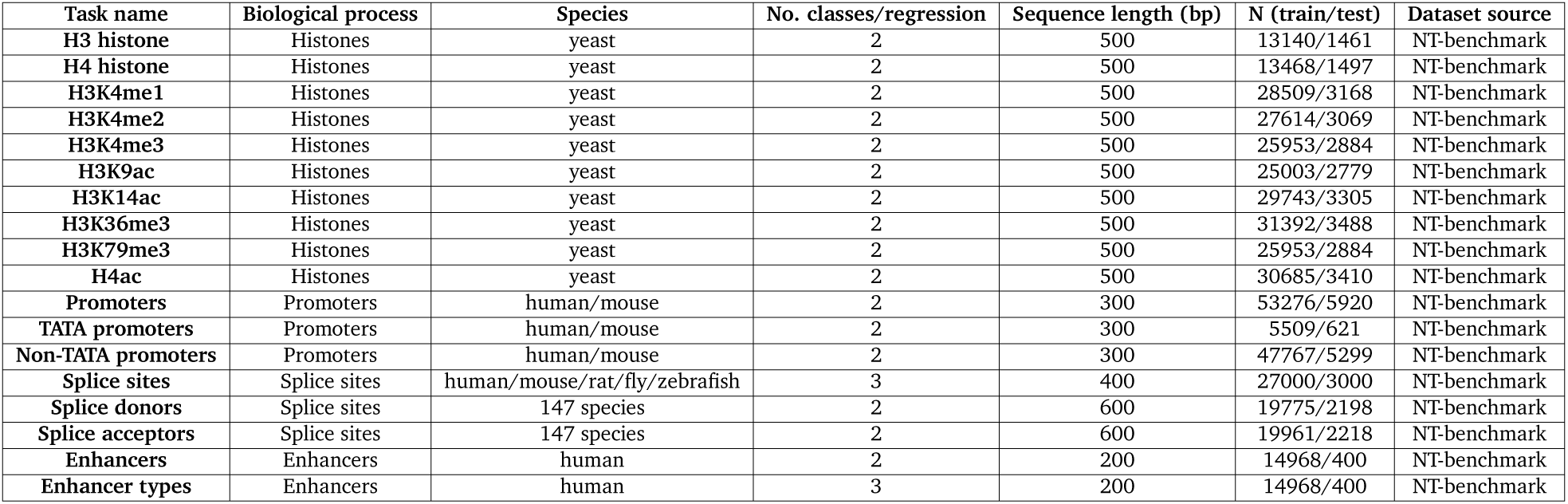
Information about all tasks in the Nucleotide Transformer benchmark[26].

| Task name | Biological process | Species | No. classes/regression | Sequence length (bp) | N (train/test) | Dataset source |
| --- | --- | --- | --- | --- | --- | --- |
| H3 histone | Histones | yeast | 2 | 500 | 13140/1461 | NT-benchmark |
| H4 histone | Histones | yeast | 2 | 500 | 13468/1497 | NT-benchmark |
| H3K4me1 | Histones | yeast | 2 | 500 | 28509/3168 | NT-benchmark |
| H3K4me2 | Histones | yeast | 2 | 500 | 27614/3069 | NT-benchmark |
| H3K4me3 | Histones | yeast | 2 | 500 | 25953/2884 | NT-benchmark |
| H3K9ac | Histones | yeast | 2 | 500 | 25003/2779 | NT-benchmark |
| H3K14ac | Histones | yeast | 2 | 500 | 29743/3305 | NT-benchmark |
| H3K36me3 | Histones | yeast | 2 | 500 | 31392/3488 | NT-benchmark |
| H3K79me3 | Histones | yeast | 2 | 500 | 25953/2884 | NT-benchmark |
| H4ac | Histones | yeast | 2 | 500 | 30685/3410 | NT-benchmark |
| Promoters | Promoters | human/mouse | 2 | 300 | 53276/5920 | NT-benchmark |
| TATA promoters | Promoters | human/mouse | 2 | 300 | 5509/621 | NT-benchmark |
| Non-TATA promoters | Promoters | human/mouse | 2 | 300 | 47767/5299 | NT-benchmark |
| Splice sites | Splice sites | human/mouse/rat/fly/zebrafish | 3 | 400 | 27000/3000 | NT-benchmark |
| Splice donors | Splice sites | 147 species | 2 | 600 | 19775/2198 | NT-benchmark |
| Splice acceptors | Splice sites | 147 species | 2 | 600 | 19961/2218 | NT-benchmark |
| Enhancers | Enhancers | human | 2 | 200 | 14968/400 | NT-benchmark |
| Enhancer types | Enhancers | human | 3 | 200 | 14968/400 | NT-benchmark |

**Supplementary Table 2.**
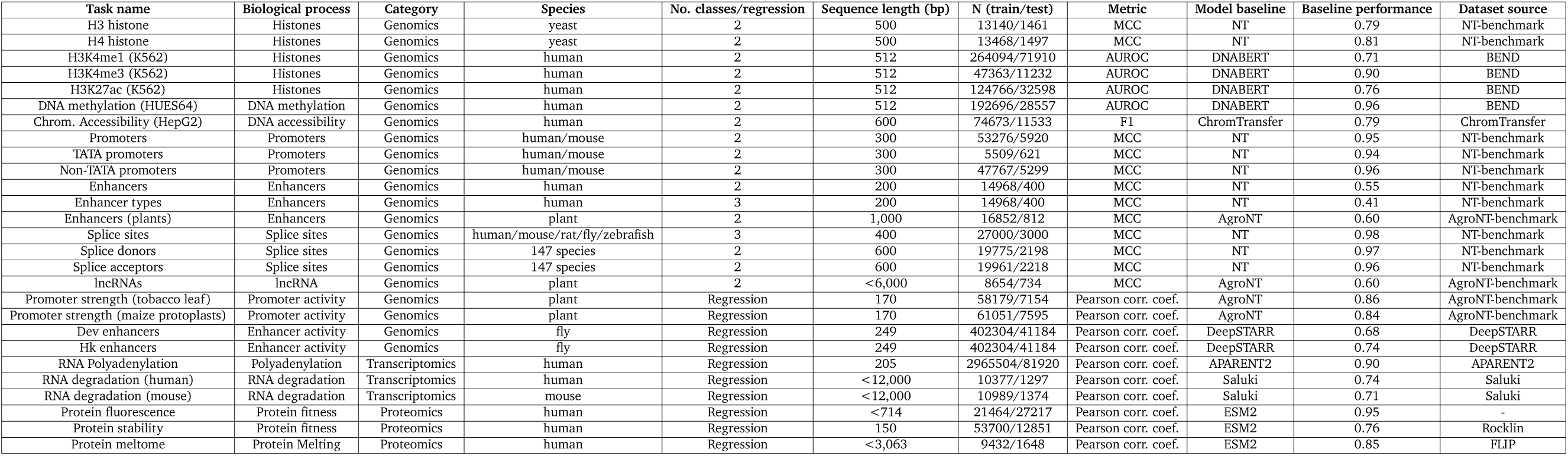
Information about all tasks and respective baseline performance and metrics.

## Supplementary Figures

**Supplementary Figure 1.**
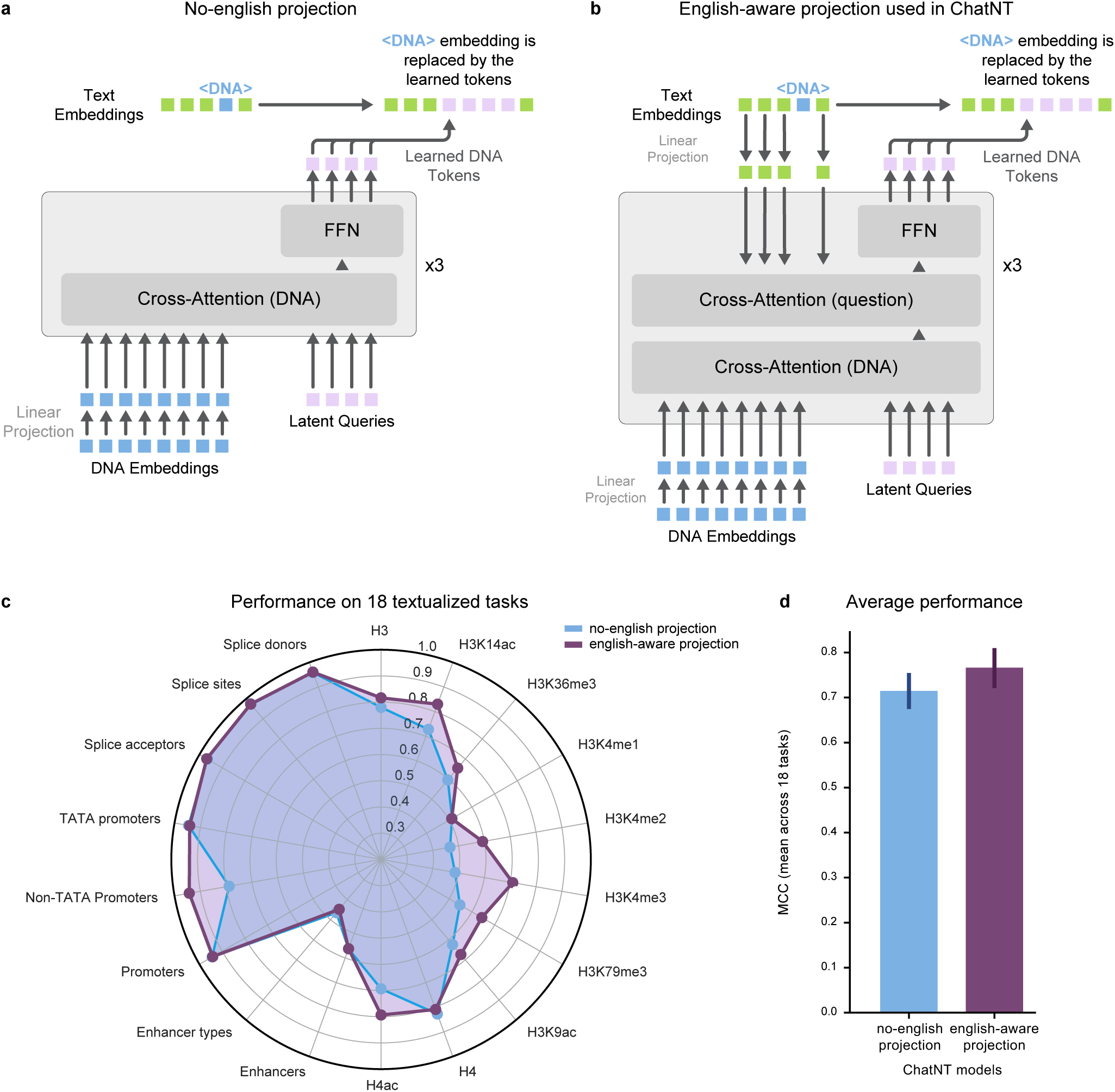
Perceiver projection. **a)** Projection without cross-attention to English question. **b)** Projection with cross-attention to english question used in final ChatNT. **c)** Radar plot comparing Chat-NT with the two projections on the 18 textualized tasks of the Nucleotide Transformer benchmark [26]. MCC performance per task is shown. **d)** Average performance across the 18 tasks. Bar-plots display the average MCC over all tasks and the standard error of the mean.

**Supplementary Figure 2.**
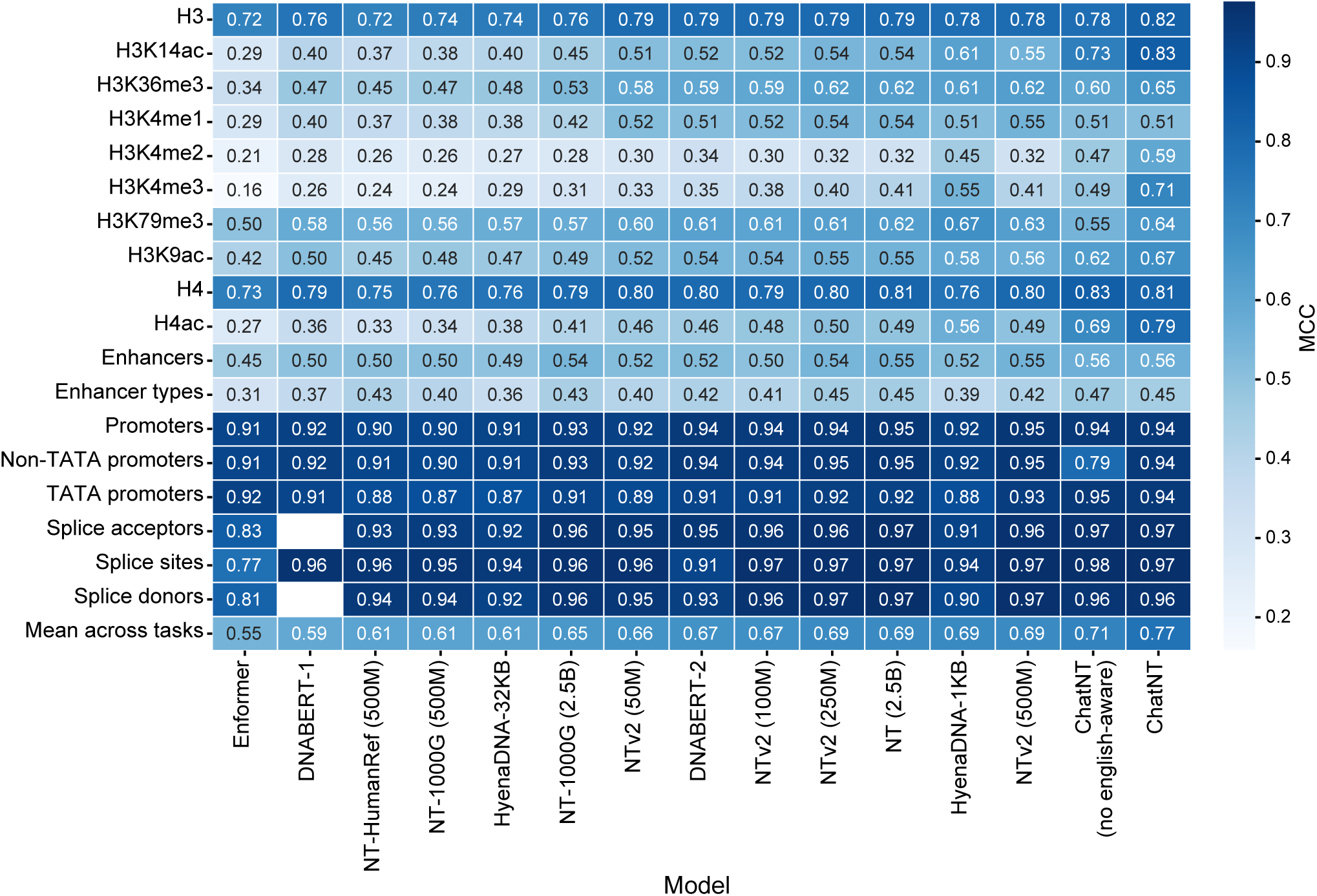
Performance of ChatNT, ChatNT with no english-aware projection and 13 different foundation models on the 18 tasks from the Nucleotide Transformer benchmark. [26].

**Supplementary Figure 3.**
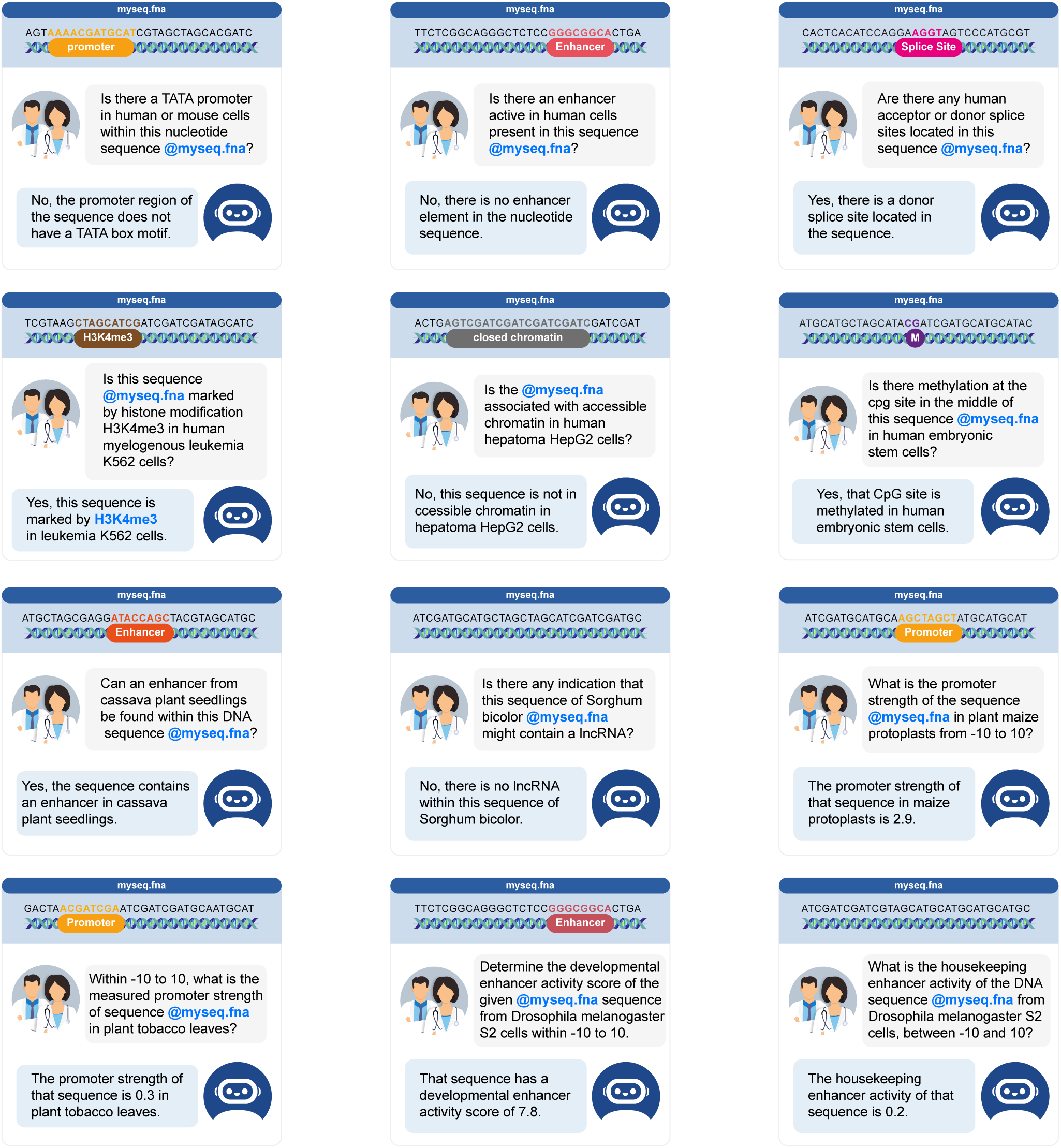
Examples of conversations included in ChatNT training data for different genomics tasks.

**Supplementary Figure 4.**
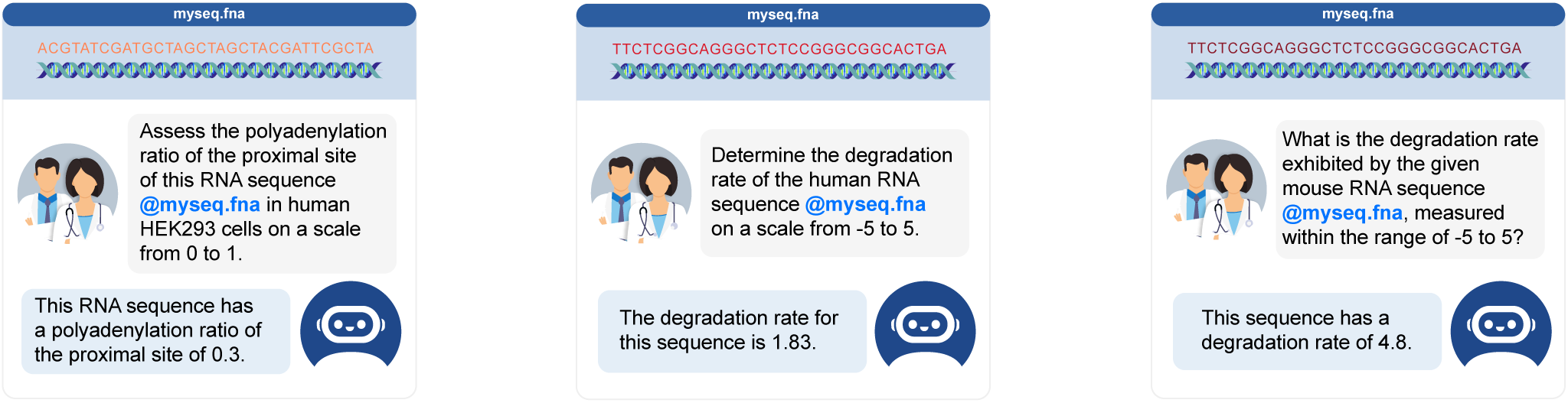
Examples of conversations included in ChatNT training data for different RNA tasks, using the respective complementary DNA sequence.

**Supplementary Figure 5.**
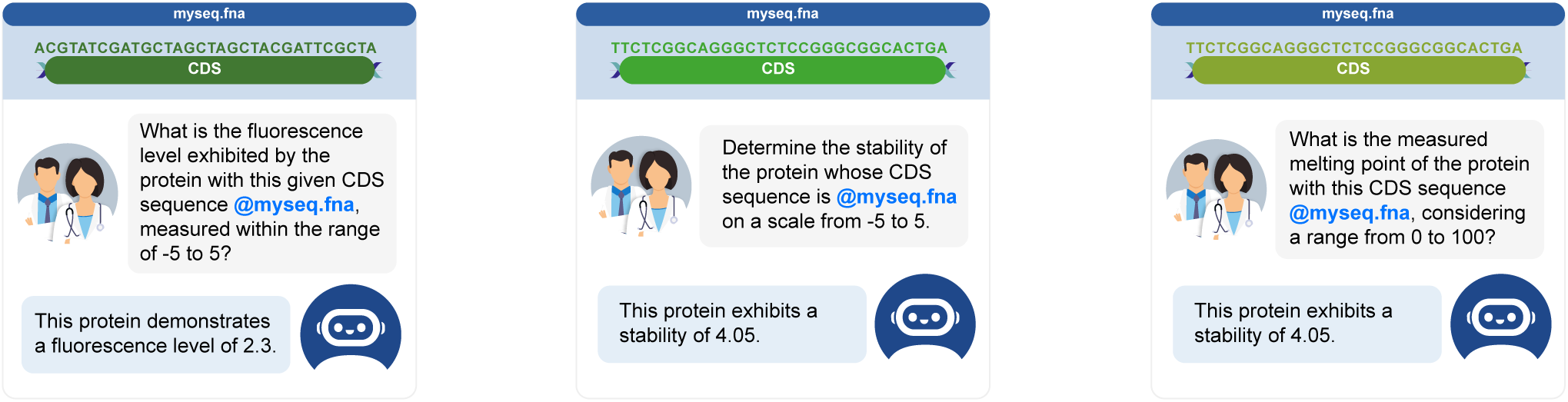
Examples of conversations included in ChatNT training data for different protein tasks, using the respective CDS sequence.

### Supplementary Pseudocode

The provided pseudocode is intended for illustrative purposes only and is not meant to be executable. It outlines the algorithmic steps and key concepts underlying our multi-modal model. In our implementation, we utilized JAX to leverage its high-performance computing capabilities and optimization techniques. In the pseudocode below we make the following assumptions:

- We use a Nucleotide Transformer model to transform DNA sequences into embeddings. Here the *nucleotide_transformer* function returns the final embedding vector before the language model head.
- We use a Llama model to decode prompt embeddings into answers. Here the *llama* object can be used either to directly generate a sentence given a prompt embedding or to compute perplexity over a candidate answer given a prompt embedding.
- For the sake of readability and understandability we do not show code and infrastructure optimization made, though in our implementation we relied on data parallelism, model parallelism, mixed precision, gradient accumulation and gradient checkpointing.
- We present in the code below the English-aware projection module that led to the best results during our experiments. This module uses a multi-modal perceiver resampler architecture and has two main goals: (1) projecting the DNA embeddings produced by the Nucleotide Transformer into the input embedding space of Llama and (2) resample the DNA embeddings into a fixed size embedding that does not depend anymore of the DNA sequence length.
- We present how we parse the model’s output to compute evaluation metrics for regression and cliassification tasks. In both cases, our code makes the assumption that the model’s output is probably structured and that the label / floating value can be extracted directly with a regex. While there is no constraint enforcing this in practice and that the model could produce any arbritary answer, we observed in our experiments that after a few gradient descents only, the model outputs structured answers satisfying these constraints. We haven’t observed counter examples so far in our experiments. However, this script wouldn’t work to evaluate a randomly initialized model.

---

**Supplementary Pseudocode 1.**
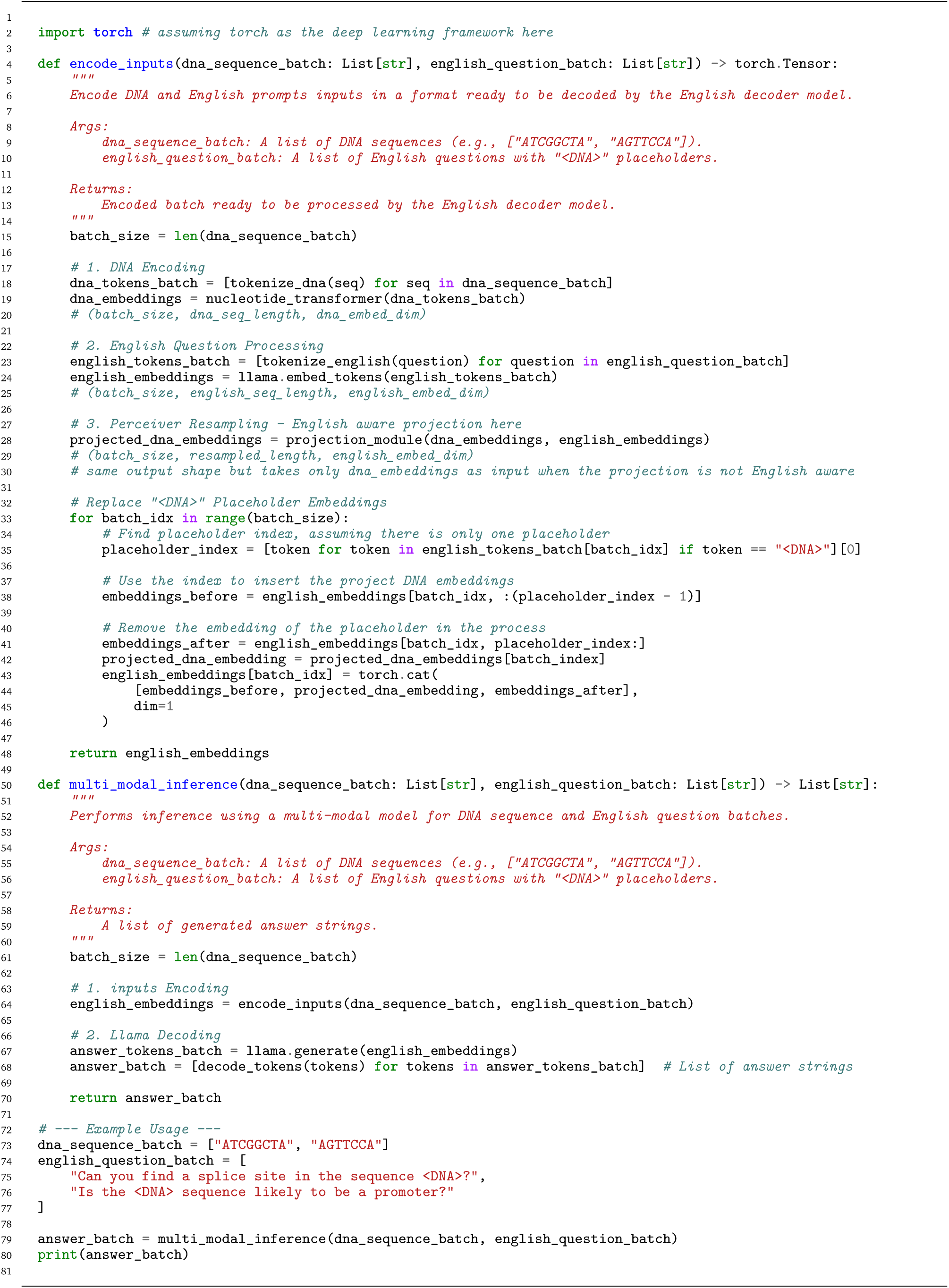
Multi-Modal Inference.

**Supplementary Pseudocode 2.**
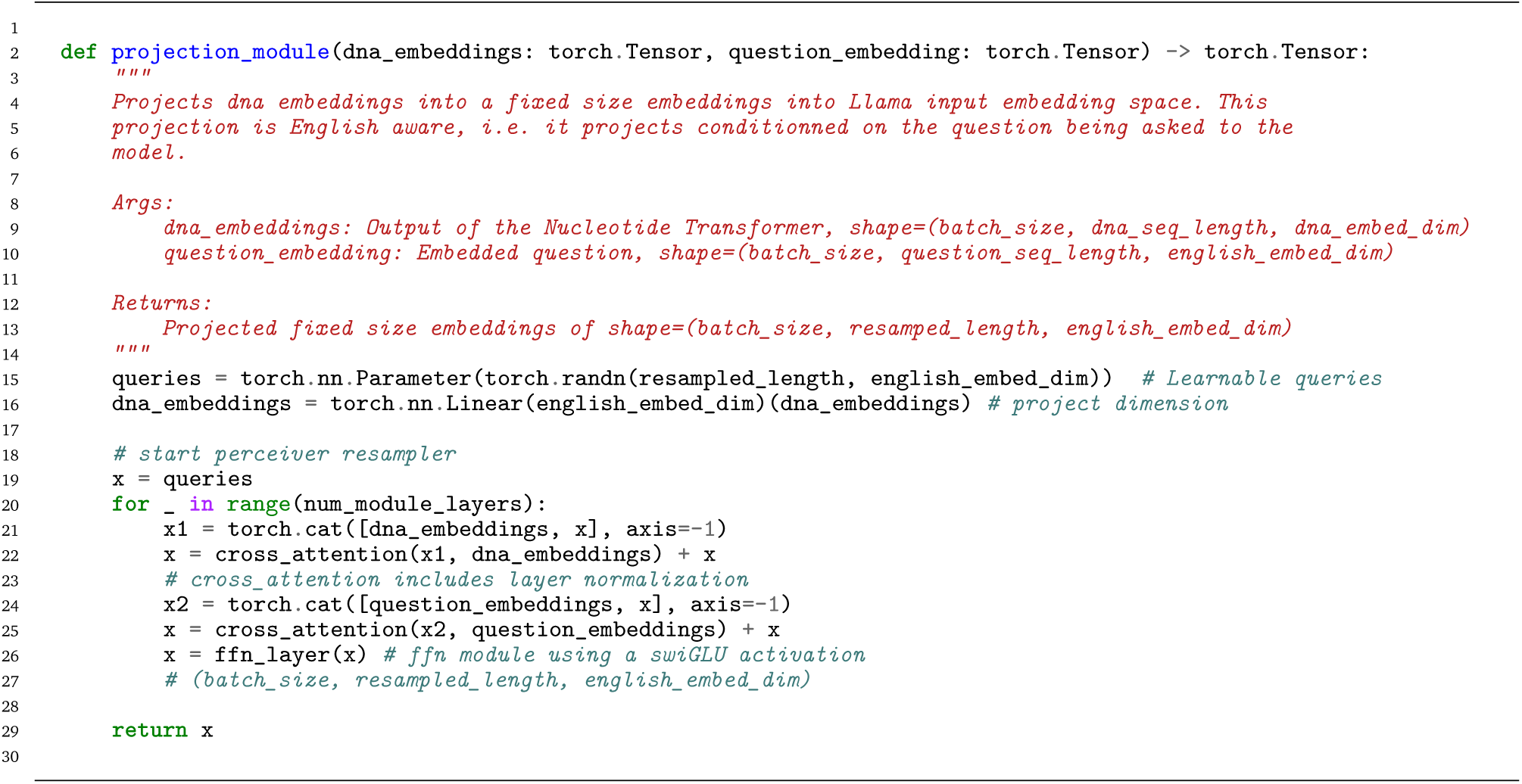
English-aware Projection Module.

**Supplementary Pseudocode 3.**
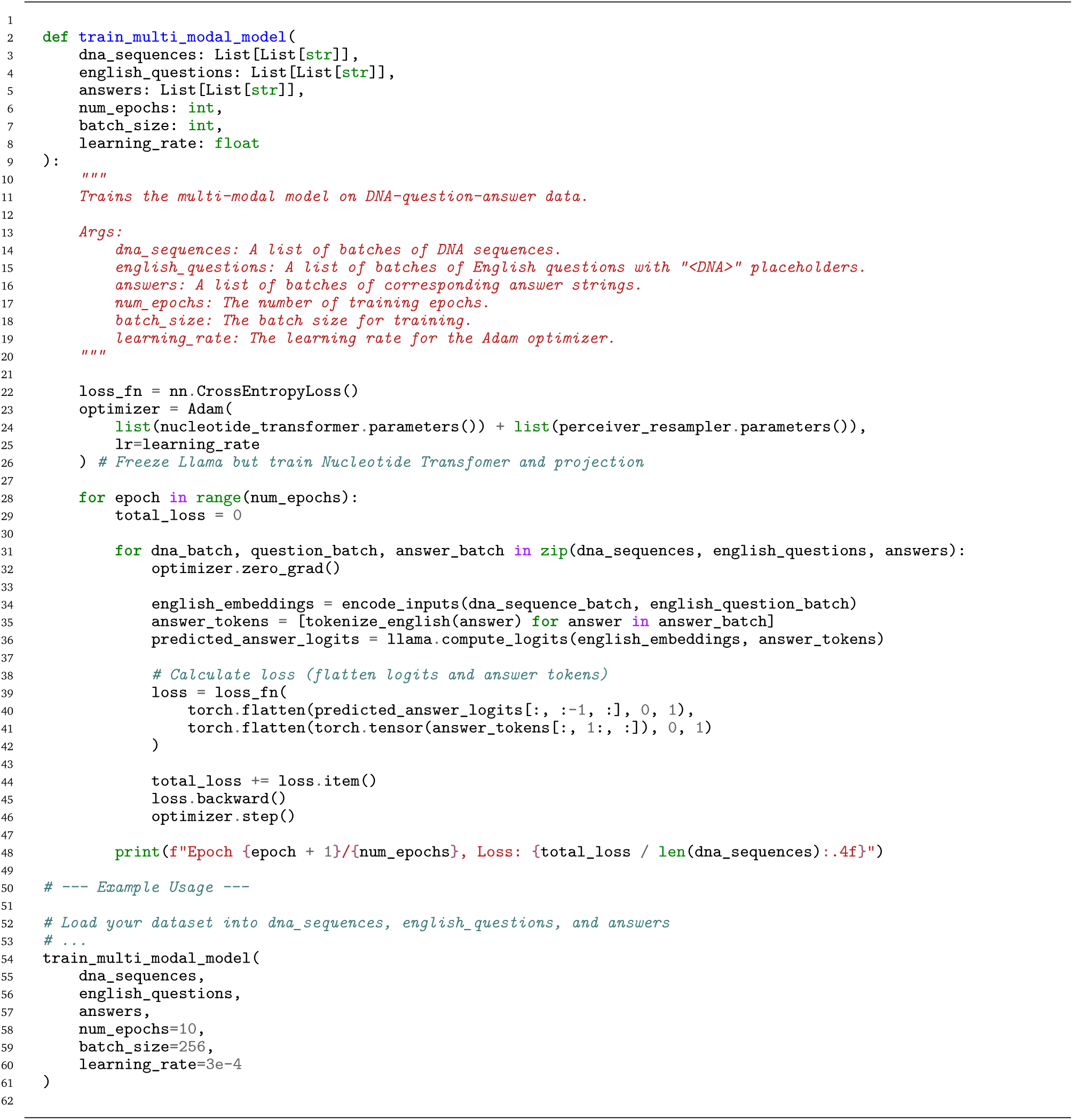
Multi-Modal Training.

**Supplementary Pseudocode 4.**
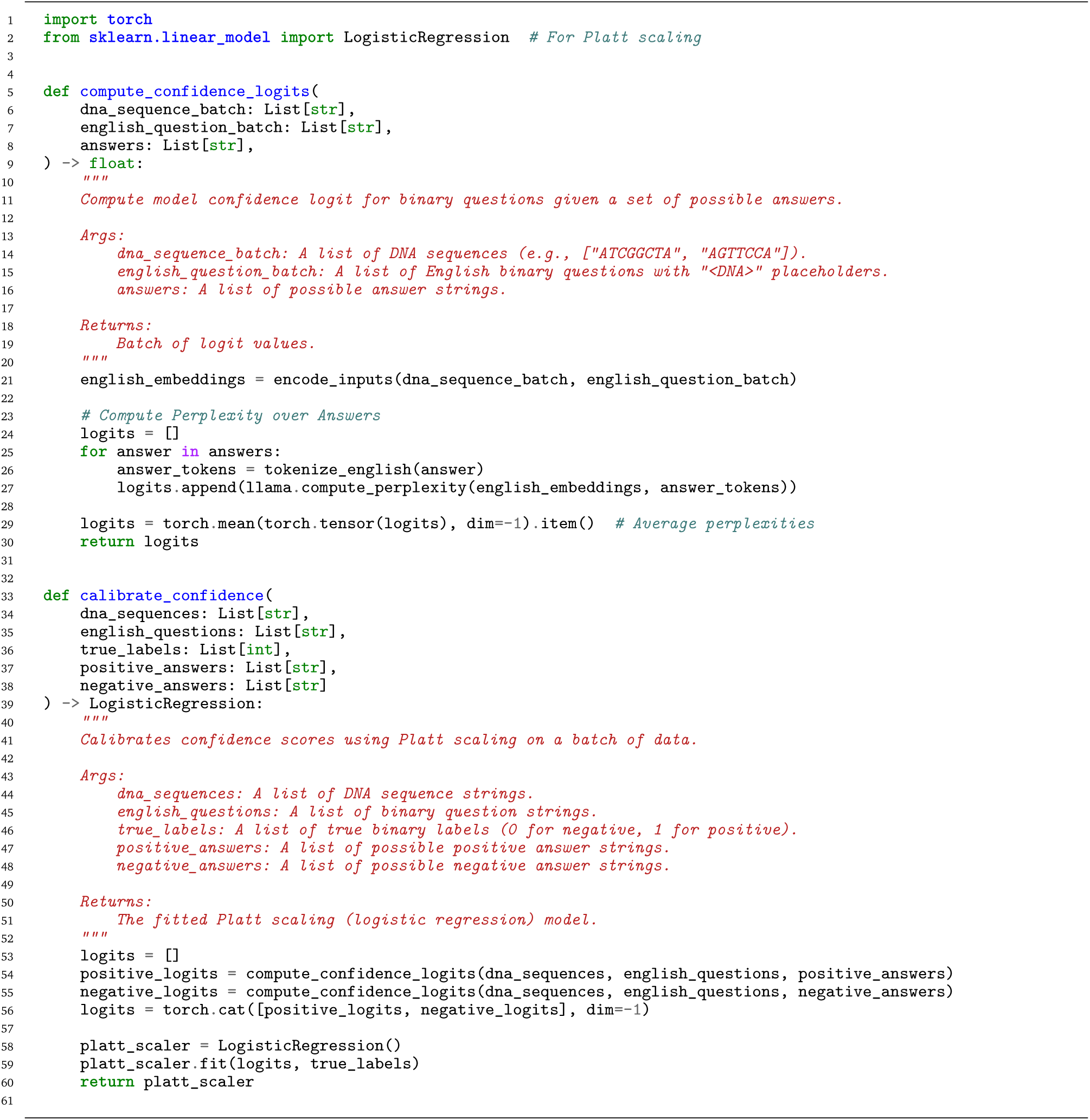
Evaluation and calibration of model’s confidence on binary classification tasks.

**Supplementary Pseudocode 5.**
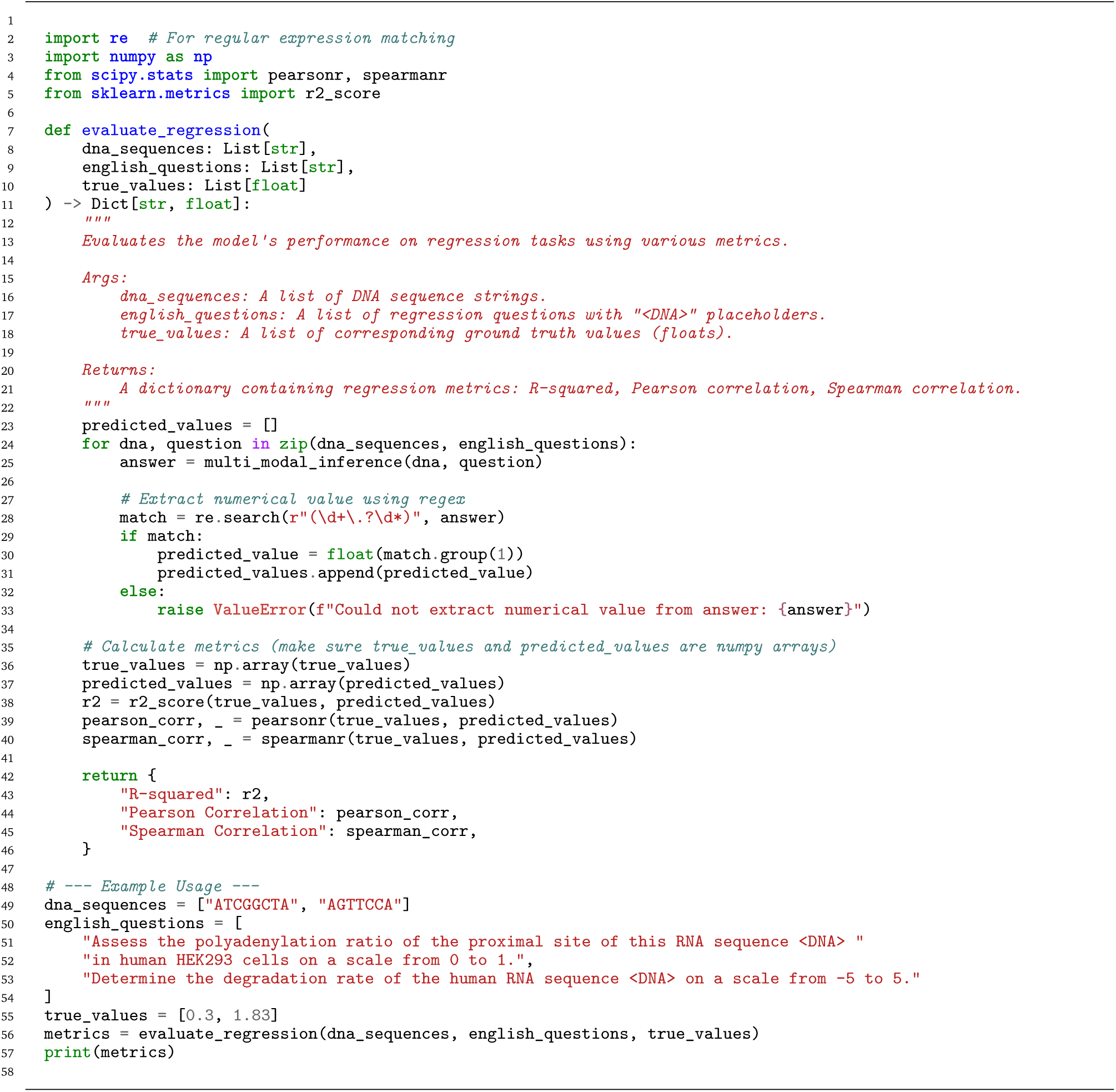
Model’s evaluation for regression tasks.

**Supplementary Pseudocode 6.**
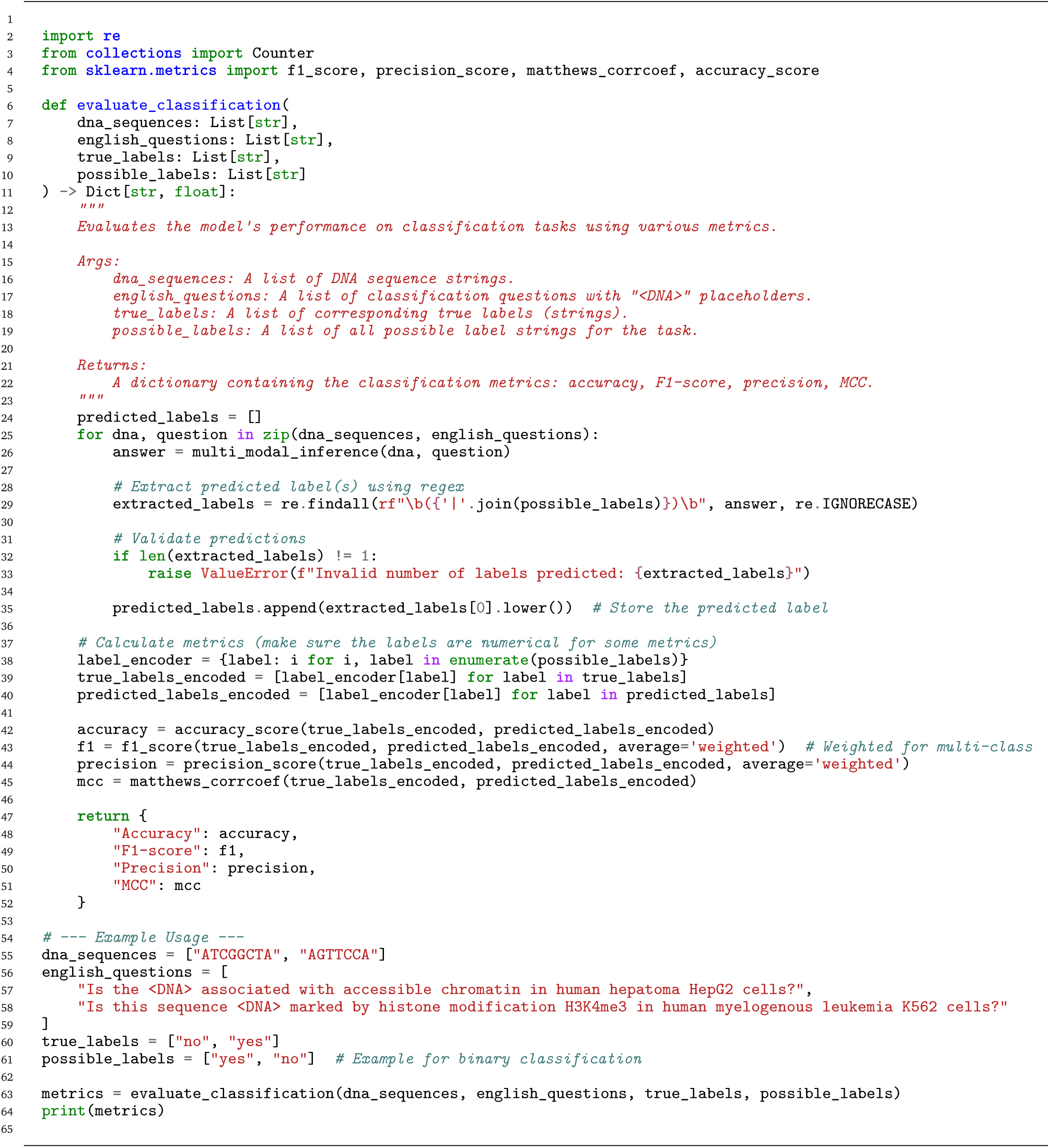
Model’s evaluation for classification tasks.

## Notes

### Summary of Updates

We have added pseudo-code in the supplementary material.

